# Less is More: Nek2A Unclusters Extra Centrosomes and Induces Cell Death in Cancer Cells via KIF2C Interaction

**DOI:** 10.1101/2023.11.21.567992

**Authors:** Batuhan Mert Kalkan, Selahattin Can Özcan, Enes Çiçek, Mehmet Gönen, Ceyda Açılan

## Abstract

Unlike normal cells, cancer cells frequently exhibit extra centrosomes, leading to formation of multipolar spindles that can trigger cell death. Nevertheless, they manage to divide successfully and escape the deadly consequences of unequal segregation of genomic material by coalescing their extra centrosomes into two poles. This unique trait of cancer cells presents a promising target for cancer therapy, focusing on selectively attacking cells with supernumerary centrosomes.

Nek2A is a kinase involved in mitotic regulation, including the centrosome cycle, where it phosphorylates linker proteins to separate centrosomes. In this study, we investigated if Nek2A also unclusters extra centrosomes, akin to its separation function. Reduction of Nek2A activity, achieved through knockout, silencing, or inhibition, promotes centrosome clustering, whereas its overexpression results in unclustering. Significantly, this unclustering activity induces cell death, but only in cancer cells with extra centrosomes, both *in vitro and in vivo*. Notably, none of the known centrosomal (e.g., CNAP1, Rootletin, Gas2L1) or non-centrosomal (e.g., TRF1, HEC1) Nek2A targets were implicated in this unclustering activity. Additionally, Nek2A operated via a mechanism distinct from other unclustering factors like HSET and NuMA.

Through TurboID proximity labeling analysis, we identified novel proteins associated with the centrosome or microtubules, expanding the known interaction partners of Nek2A. KIF2C, in particular, emerged as a novel interactor, confirmed through coimmunoprecipitation and localization analysis. The silencing of KIF2C diminished the impact of Nek2A on centrosome unclustering and rescued cell viability. Additionally, elevated Nek2A levels were indicative of better patient outcomes, specifically in those predicted to have excess centrosomes. Therefore, while Nek2A is a proposed target, its use must be specifically adapted to the broader cellular context, especially considering centrosome amplification. Discovering partners such as KIF2C offers fresh insights into cancer biology and new possibilities for targeted treatment.

## INTRODUCTION

Centrosomes are key to microtubule organization and cell division, with their dysfunction linked to diseases like cancer, where cells often have excess centrosomes that promote genomic instability and tumor growth [1-3]. For cancer cells with extra centrosomes, clustering is crucial to achieve bipolar division and sustain their survival. This process enables the formation of a functional bipolar spindle in mitosis, averting the emergence of lethal multipolar spindles. Hence, understanding the processes that govern centrosome clustering is vital to pinpoint new therapeutic strategies aimed at disrupting this clustering, offering a selective means to eliminate cancer cells.

One of the major regulators that control the centrosome cycle and separation is Nek2A kinase [4,5]. Nek2A orchestrates centrosome separation by phosphorylating centrosomal linker proteins, such as C-Nap1 and rootletin, leading to their disassembly at the onset of mitosis [6-8]. This phosphorylation event weakens the cohesion between centrosomes, allowing them to move apart and form the poles of the mitotic spindle, which is crucial for proper chromosome segregation. Although the role of Nek2A in centrosome separation is well-documented, it was unclear if similar mechanisms apply to the disjunction of supernumerary centrosomes and if Nek2A could facilitate their unclustering.

In this study, we investigated the potential of Nek2A to regulate unclustering of centrosomes in cancer cells with extra centrosomes, and whether impeding Nek2A-mediated unclustering could emerge as a potential cancer treatment strategy. Our findings showed that while Nek2A overexpression could indeed uncluster centrosomes, none of its known targets appeared to be involved in this process. Through proximity labeling, we identified a novel collaboration between Nek2A and KIF2C, a kinesin family member, that orchestrates centrosome unclustering through a mechanism independent from those described earlier. Specifically, we found that KIF2C is indispensable for the unclustering effect triggered by Nek2A, a phenomenon exclusive to cancer cells with additional centrosomes. We demonstrated that targeting cells with amplified centrosomes and elevated Nek2A/KIF2C expression could selectively kill cancer cells, a finding substantiated both *in vitro* and *in vivo*. Moreover, in support of our hypothesis, high levels of Nek2A correlated with improved patient outcomes when centrosome amplification was predicted to be high.

## RESULTS

### Nek2A Regulates Centrosome Clustering in Cancer Cells with Supernumerary Centrosomes

To evaluate Nek2A’s role in centrosome clustering regulation, we generated a doxycycline (dox)-inducible lentiviral overexpression system. This was introduced into the N1E-115 mouse neuroblastoma cell line, known for its widespread supernumerary centrosomes and effective centrosome clustering (**Supp. Figure 1A**), making it an ideal model for study [9]. We stained the cells with DAPI, γ-Tubulin and α-Tubulin to determine the number of spindle poles and distinguish bipolar clustered and multipolar metaphases based on DNA shape and planar arrangement of centrosomes (**Fig. 1A**). Overexpression of Nek2A dramatically reduced centrosome clustering in N1E-115 cells, promoting formation of multipolar spindles (MPS) during metaphase (**Fig. 1B**, p<0.01). In line with the hypothesis that centrosome clustering is vital for cancer cell survival, overexpression of Nek2A led to a marked reduction in cell viability in N1E-115 cells (p<0.001). To determine if this observation isn’t just specific to a particular cell line, we examined various human cancer cells, such as pancreatic ductal adenocarcinoma (PDAC), known for exhibiting centrosome amplification (CA) [10]. We characterised three PDAC cell lines (MIA PaCa-2, Panc1, and SU86.86) for PLK4 and STIL expressions, key indicators of CA (**Supp. Figure 1B**). Among three, SU86.86 showed the highest PLK4 levels and CA (∼22%), confirmed by γ-Tubulin staining (**Supp. Figure 1C and D**). Hence, we generated a dox-inducible Nek2A overexpression in SU86.86 cells (**Supp. Figure 1E**) similar to N1E-115 and found that Nek2A significantly promoted MPS formation (**Fig. 1C**) and reduced cell viability (**Fig. 1D**), further implicating its role in centrosome unclustering which trigger cell death.

**Figure 1:**
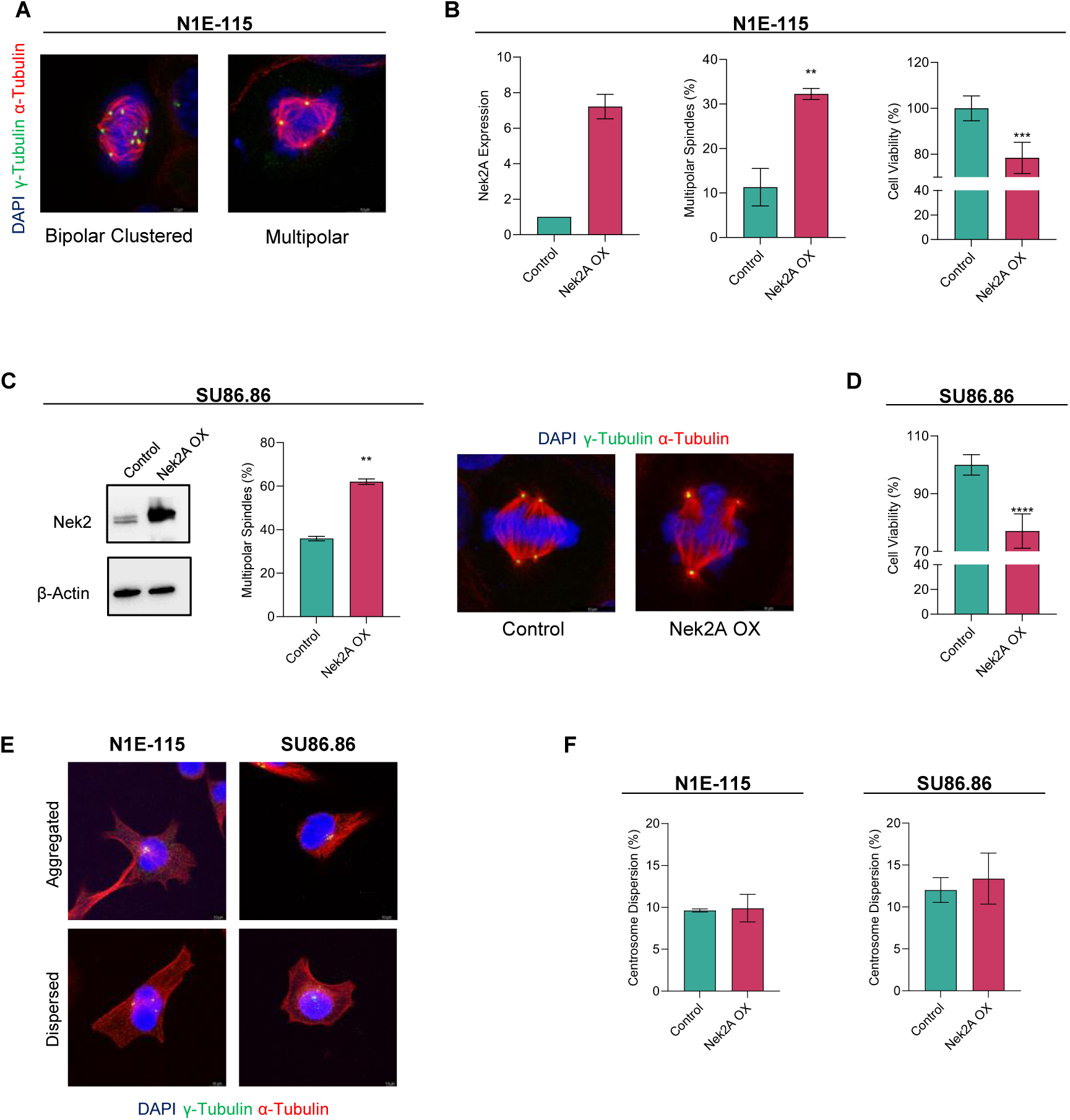
Nek2 overexpression reduces centrosome clustering in cancer cells naturally comprising supernumerary centrosomes. (**A**) Representative IF staining images demonstrating supernumerary centrosomes, bipolar-clustered and multipolar metaphases observed in N1E-115 cells. (**B**) Ectopic overexpression of Nek2A confirmed by Western Blot (left), metaphase scoring indicating the induction of MPS formation (middle) and cell viability assay (right) in N1E-115 cells. (**C**) Overexpression of Nek2A confirmed by RT-qPCR (left), metaphase scoring (middle) and representative IF staining images (right) showing bipolar with clustered centrosomes and multipolar metaphases in SU86.86 cells. (**D**) WST-1 assay showing significant decrease in viability of SU86.86 cells when Nek2A is overexpressed for 72 hours. (**E**) Representative images demonstrating aggregated and dispersed localization of supernumerary centrosomes in N1E-115 and SU86.86 cells. All experiments were performed as two biological repeats with at least 200 metaphases with CA scored per experiment. Error bars represent standard deviation from the mean. Statistical significance was shown as * : p<0.05, **: p<0.01, ***: p<0.001, **** : p<0.0001. OX: Overexpression

Interestingly, during interphase, extra centrosomes often appeared clustered together (**Fig. 1E**). Thus, we explored if Nek2A overexpression, which we found affects centrosome coalescence during mitosis, could also disperse these centrosomes in interphase. However, Nek2A did not alter the arrangement of additional centrosomes in either N1E-115 or SU86.86 cells (**Fig. 1F**), suggesting that Nek2A’s unclustering effect is mitosis-dependent.

In order to test whether Nek2A can still regulate centrosome clustering in cells without amplification or low levels of CA, we overexpressed Nek2A in MDA-MB-231 (∼7% CA) and U2OS cells (1% CA) (**Supp. Fig. 1F, 1G**). Unlike cells with high CA and effective clustering, Nek2A overexpression did not promote centrosome unclustering in these cells (**Supp. Fig. 1H**). This suggests that Nek2A’s unclustering effect is only apparent in cells with supernumerary centrosomes that are already clustered. To test this hypothesis and assess the impact of Nek2A as a possible controller of centrosome clustering, we artificially increased the number of centrosomes through nocodazole treatment and PLK4 overexpression. Indeed, CA was successfully induced following these methods (**Fig. 2A**) in U2OS and MDA-MB-231 cells. Overexpressing PLK4 for 72 hours caused CA in about 40% of MDA-MB-231 and U2OS cell populations (**Fig. 2B**). Additionally, we induced CA in U2OS cells using the cytokinesis inhibitor DCB and STILL overexpression (**Supp. Fig. 2A-B**). Post-amplification, metaphases were categorized as either bipolar clustered or multipolar. As observed in N1E-115 and SU86.86, Nek2A overexpression led to multipolar metaphases and significantly reduced centrosome clustering in MDA-MB-231 (**Fig. 2C**) and U2OS cells (**Fig. 2D**). In parallel with these results, overexpression of Nek2A induced significant increase in MPS formation in both of the alternative CA models, DCB and STIL overexpression respectively (**Supp. Fig. 2C**). To investigate whether inference with Nek2A activity exerts the reverse effects, several approaches were undertaken including knockout, RNAi and using chemical inhibitor (JH295) targeting Nek2A in two different cell lines with CA induced with the aforementioned methods (**Supp. Fig. 2D**). Consistent with our expectations, the results showed that suppression of Nek2A significantly decreased the percentage of MPS. To address whether the kinase activity of Nek2A is important for centrosome clustering, we overexpressed kinase-deficient mutant form of Nek2A (K37R substitution) [11], providing a dominant-negative (DN) phenotype. Over-expression of the DN mutant acted similarly to chemical and transcriptional inhibition of Nek2A, favouring centrosome clustering. Taken together, our data strongly argues that Nek2A regulates centrosome clustering which requires its kinase activity, in cancers cell with supernumerary centrosomes.

**Figure 2:**
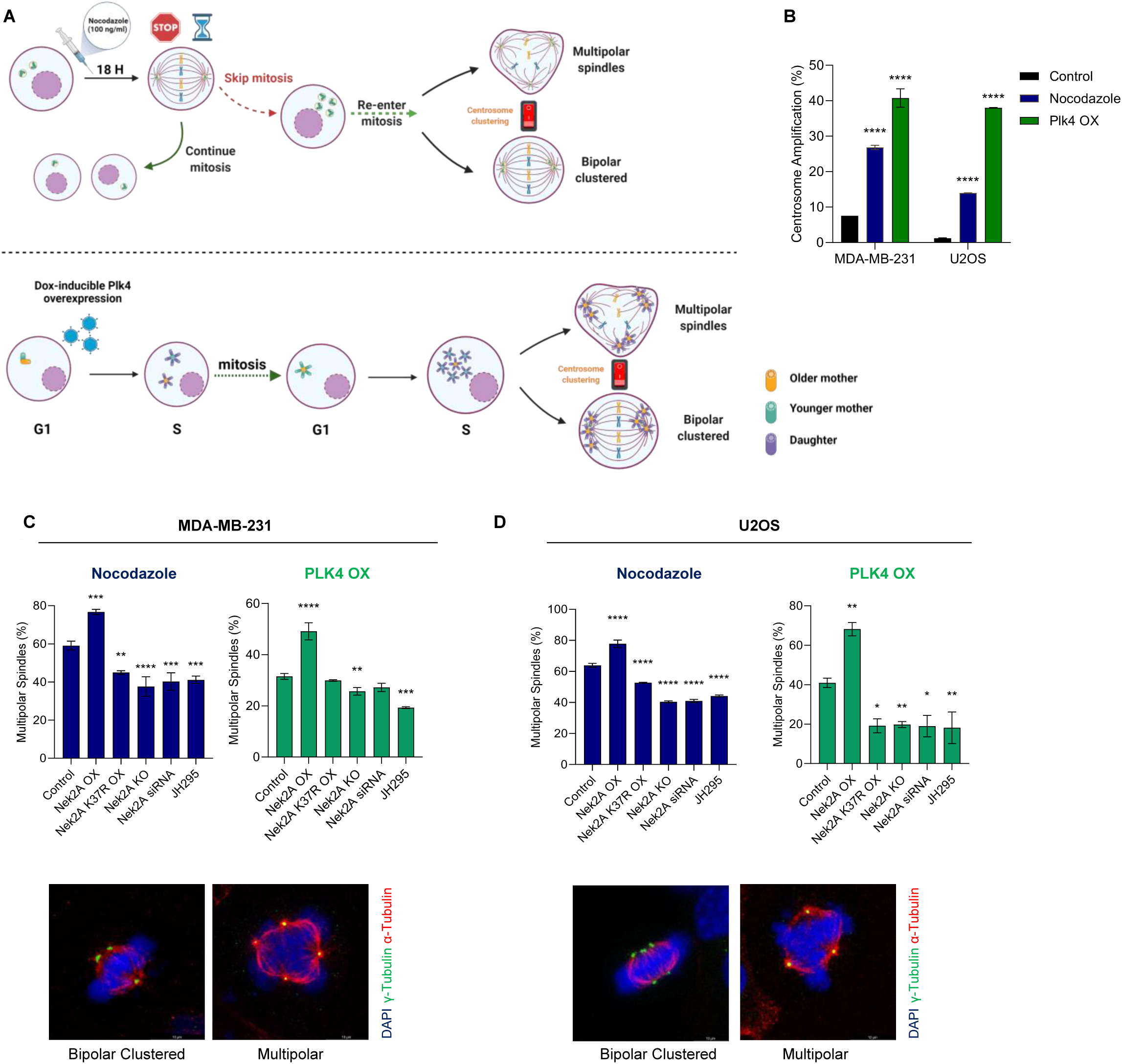
Nek2 regulates centrosome clustering in cells which are induced to have amplified centrosomes. (**A**) Experimental setups to induce centrosome amplification by nocodazole (upper panel) and overexpression of PLK4 overexpression (lower panel). Created with BioRender. (**B**) Centrosome amplification levels obtained by nocodazole and PLK4 overexpression models. (**C**) Percentage multipolarity of MDA-MB-231 cells in nocodazole (top-left), PLK4 (top-right) models and representative images (bottom) showing bipolar clustered and multipolar metaphases. (**D**) Percentage multipolarity of U2OS cells in nocodazole (top-left), PLK4 (top-right) models and representative images (bottom) showing bipolar clustered and multipolar metaphases. Experiments were performed as two biological repeats with at least 200 metaphases with CA scored per experiment. Error bars show standard deviations. Statistical significance was shown as * : p<0.05, **: p<0.01, ***: p<0.001, **** : p<0.0001. OX: overexpression, KO: knock-out,

### Nek2A Overexpression Promotes the Depletion of Cells with CA

Considering Nek2A overexpression’s effect on centrosome clustering, we performed an *in vitro* competition assay to show its negative impact on cells with CA. We engineered U2OS and MDA-MB-231 cells for dox-inducible Nek2A and PLK4 overexpression (**Supp. Fig. 3A**), labeling PLK4 and Nek2A co-expressors (Dox-Plk4&Nek2A) with H2B-mCherry and PLK4 expressors (Dox-Plk4) with H2B-eGFP (**Fig. 3A)**. Co-culturing these cells with/without doxycycline for 10 days, we noted a marked decrease in mCherry-tagged cells, undergoing multipolar divisions from Nek2A overexpression, corroborated by flow cytometry and live-cell imaging (**Fig. 3B**, **Supp. Fig. 3B**). We repeated this with MDA-MB-231 cells (**Supp. Fig. 3C**) and SU86.86 cells using SU86.86(WT)-H2B-eGFP and SU86.86(dox-Nek2A)-H2B-mCherry (**Supp. Fig. 3D**), observing similar effects. Control groups showed comparable proliferation rates. This indicates Nek2A overexpression’s detrimental role in cells with supernumerary centrosomes, leading to cell death through multipolar divisions.

**Figure 3:**
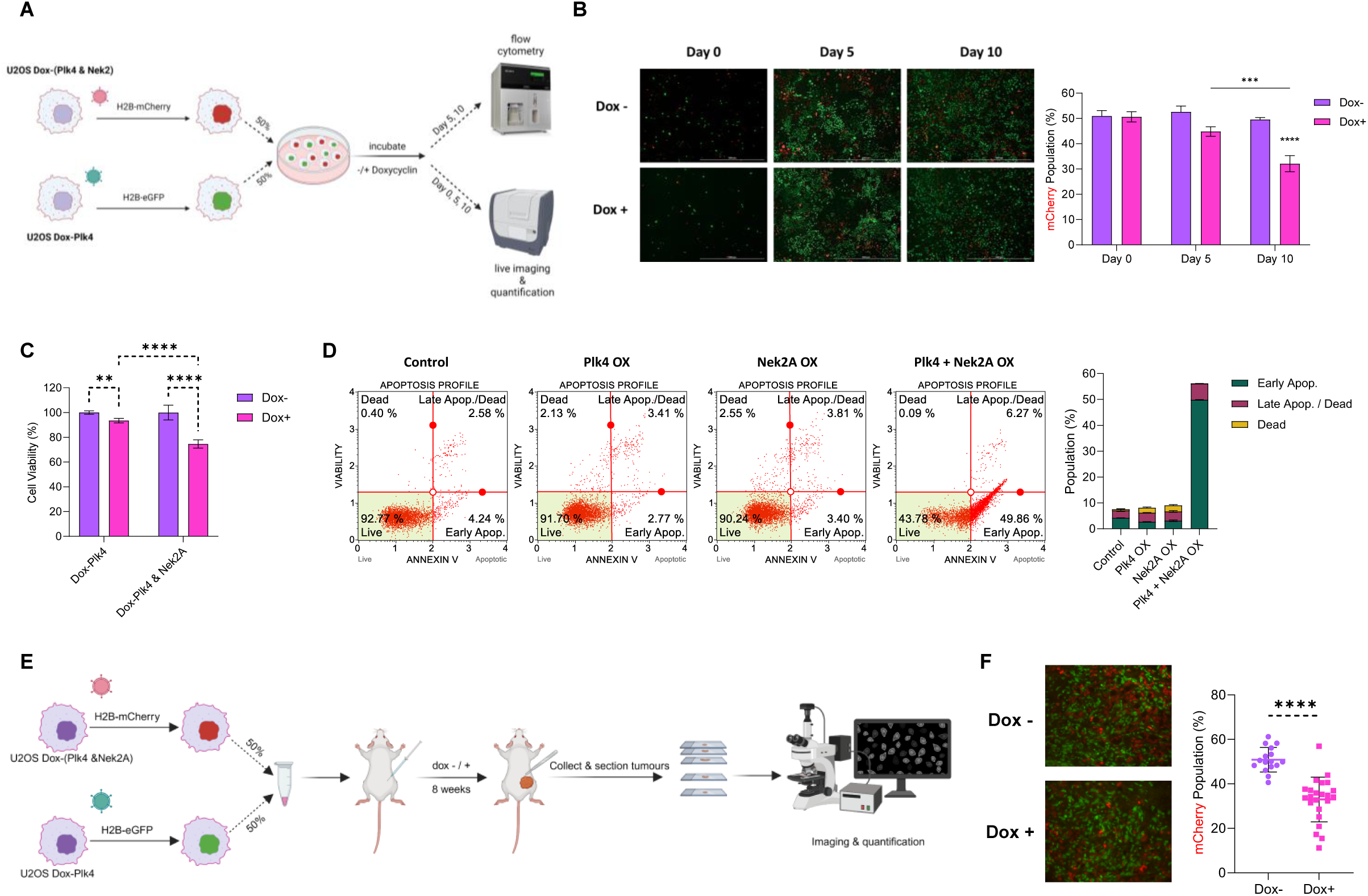
Nek2A overexpression induces depletion of cells with centrosome amplification *in vitro* and *in vivo*. (**A**) Experimental setup of *in vitro* competition assay. Image created with BioRender. (**B**) Imaging-based analysis and quantification of mCherry-tagged U2OS cells co-overexpressing Plk4 and Nek2A. (**C**) WST-1 assay showing the significant decrease in cell viability as a result of Nek2A overexpression in cells with CA (induced by Plk4 overexpression) (**D**) Annexin V assay confirming the apoptosis induced by multipolar metaphases. (**E**) Experimental setup of *in vivo* competition assay. (**F**) Representative images of tumour tissue slices and quantification of mCherry-tagged U2OS Dox (PLK4 & Nek2) cells. Experiments were performed as at least 2 biological replicates. Error bars show standard deviations. Statistical significance was shown as * : p<0.05, **: p<0.01, ***: p<0.001, **** : p<0.0001. OX: overexpression

To ascertain whether the decrease in cell viability in these cells stemmed from programmed cell death pathways or a reduction in cell division, we assessed the population of cells positive for Annexin V (**Fig. 3D**) and examined Caspase 3/7 activity (**Supp. Fig. 3E**) in cells that either overexpressed Nek2A without CA, exhibited CA without Nek2A overexpression, or had both conditions simultaneously. As we had predicted, only cells with both PLK4 and Nek2A overexpression underwent apoptosis, providing strong evidence that the unclustering effect induced by Nek2A leads to the observed cell death phenotype only in the presence of extra centrosomes.

Lastly, we conducted an *in vivo* version of the competition assay by subcutaneously implanting a mix of eGFP-tagged U2OS dox(PLK4) and mCherry-tagged U2OS Dox(PLK4&Nek2A) cells (**Fig. 3E**). After eight weeks, we analyzed the tumors and found a significant reduction in Nek2A overexpressing (mCherry+) cells with CA (**Fig. 3F**). This demonstrates that Nek2A-induced multipolar divisions lead to cell depletion in both *in vitro* and *in vivo* settings.

### Nek2A Overexpression Induces MPS Independent of its Intercentriolar Linker and Chromosomal Targets

To comprehend the mechanism behind Nek2A’s unclustering activity, we initially investigated the involvement of its possible targets. Nek2A kinase, known for regulating the centrosome cycle by phosphorylating C-Nap1 (CEP250) [7,12], Rootletin (CROCC) [8,13], and GAS2L1 [14,15], also has non-centrosomal targets like Hec1 [16] and Trf1 [17] that may influence microtubule attachments and contribute to centrosome unclustering (**Fig. 4A**). We postulated that suppressing these targets would eliminate the observed phenotype resulting from Nek2A overexpression. Therefore, we generated monoclonal C-Nap1 and Rootletin knockout U2OS (dox-Nek2A) cells (**Supp. Fig. 4A-B)** and scored metaphases following induction of CA via two independent methods and Nek2A overexpression. As expected, Nek2A overexpression increased MPS and reduced bipolar clustered metaphases. Intriguingly, C-Nap1 knockout impaired centrosome clustering in both CA models, irrespective of Nek2A **(Fig. 4B)**. In the nocodazole-induced CA model, Nek2A overexpression in C-Nap1 knockout cells resulted in a further decline in centrosome clustering, suggesting that absence of C-Nap1 promotes the formation of MPS independent of Nek2A activity. However, in the Plk4 CA model, Nek2A overexpression in C-Nap1 knockouts didn’t further reduce clustered metaphases, indicating different CA mechanisms produce varied responses. These findings were confirmed with independent monoclonal C-Nap1 knockout cells (**Supp. Fig. 4C)**.

**Figure 4:**
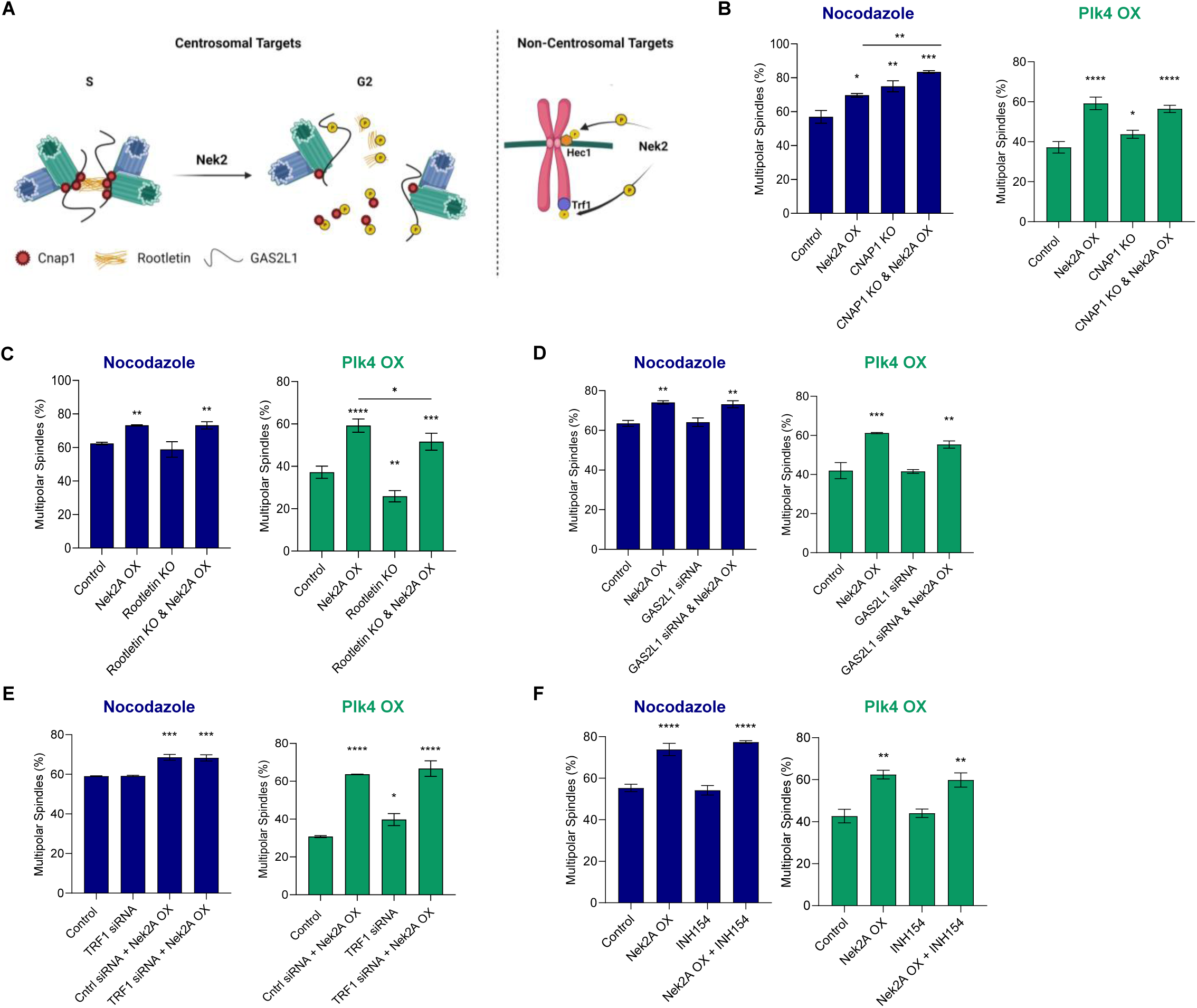
Nek2A overexpression induces MPS formation independent of its intercentriolar linker and chromosomal targets. (**A**) Known centrosomal and non-centrosomal targets of Nek2A kinase. Image created by BioRender. Percentage MPS of U2OS cells in nocodazole and PLK4 models to examine effects of (**B**) CNAP1, (**C**) Rootletin (**D**) GAS2L1, (**E**) TRF1 and (**F**) Hec1 (INH154 inhibits Nek2-Hec1 interaction). Experiments were performed as two independent repeats with at least 200 metaphases with CA scored per experiment. . Error bars show standard deviations. Statistical significance was shown as * : p<0.05, **: p<0.01, ***: p<0.001, **** : p<0.0001. OX: overexpression, KO: knock-out

To assess Rootletin’s role in Nek2-induced centrosome unclustering, we generated Rootletin knockout cells (**Supp. Fig. 4B**) and examined metaphases post Nek2 overexpression in different CA models. Rootletin’s absence reduced MPS formation in the Plk4 overexpression model but not in the nocodazole model, once again suggesting variances between the models (**Fig. 4C**). Additionally, Rootletin knockout slightly enhanced centrosome clustering in the Plk4 model, hinting at its independent role from Nek2A kinase. However, Rootletin’s absence didn’t negate Nek2A’s unclustering effect (**Fig. 4C**).

We also analyzed C-Nap1 and Rootletin knockouts’ impact on centrosome distance at interphase. As reported previously, Immunofluorescence staining with anti-γ-tubulin antibodies showed C-Nap1 loss significantly increased the centrosome distance [18,19], indicating potential dispersion-driven clustering disruption **(Supp. Fig. 4D)**. Nevertheless, Rootletin knockout didn’t notably change centrosome distance.

Lastly, we investigated GAS2L1. Despite successful GAS2L1 suppression (**Supp. Fig. 4E**), its loss didn’t affect centrosome unclustering or Nek2A-induced multipolarity any of the CA models (**Fig. 4D**).

Beyond Nek2A’s centrosomal targets, we also investigated two non-centrosomal targets, TRF1 and Hec1, for their potential roles in centrosome clustering. Using siRNA to silence TRF1 (**Supp. Fig. 4F**), we found that its suppression didn’t affect centrosome clustering in the nocodazole CA model nor did it prevent unclustering with Nek2A overexpression, even increasing multipolar metaphases in the Plk4 CA model (**Fig. 4E**). Further, Nek2A overexpression in TRF1-silenced cells still led to multipolar divisions, indicating TRF1’s non-essential role in Nek2A-induced unclustering.

For Hec1, we treated U2OS cells with INH154, disrupting Nek2A-Hec1 interaction, and observed no significant impact on centrosome clustering or reversal of Nek2A overexpression effects during metaphase (**Fig. 4F**). Thus, our findings suggest that neither TRF1 nor Hec1 are key components in Nek2A’s molecular mechanism.

### Interaction Between Nek2A and KIF2C Mediates Centrosome Unclustering

Since none of the known interactors of Nek2A appeared to be responsible for its effect on centrosome declustering, we hypothesized that a novel partner might be involved. Using the TurboID proximity labeling system and proteomic tools, we aimed to find unrecognized partners facilitating Nek2A in centrosome unclustering in cancer (**Fig. 5A**). We first generated FLAG-TurboID-Nek2A and FLAG-TurboID-Nek2A(K37R) constructs (**Supp. Fig. 5A**) and confirmed their cellular location (**Fig. 5B**), showing that both forms of Nek2A localizes to centrosomes similar to endogenous Nek2A. Additionally, streptavidin staining also confirmed that the majority of biotinylated proteins were centrosome-associated, suggesting that the identified and enriched proteins are probable participants in centrosome clustering mechanisms **(Supp. Fig. 5B)**. Our analysis confirmed known Nek2A interactors, validating our system’s reliability. It identified biotinylated Nek2A targets (CROCC & LRRC45), known centrosome clustering regulators (NuMA & KIFC1), and potential new partners like KIF2C (**Fig. 5C**). The cellular component analysis revealed that both wild-type (WT) and kinase-deficient (KD) Nek2A primarily interact with proteins found in spindles and microtubules, demonstrating similar interaction profiles for both WT and KD forms. (**Fig. 5D**).

**Figure 5:**
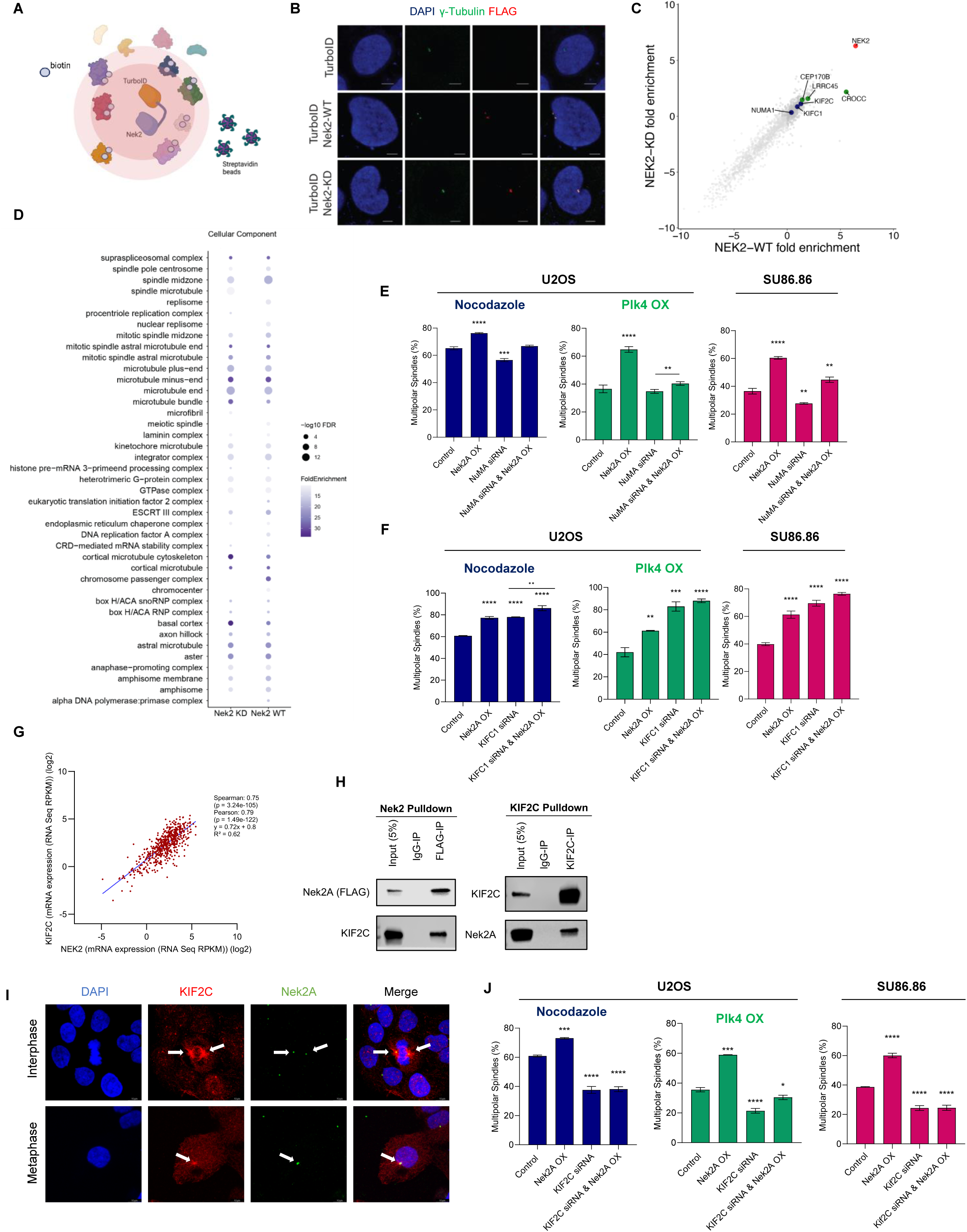
Proximity labelling and Co-IP reveals interaction between Nek2A and KIF2C regulating centrosome clustering. (**A**) Turbo-ID proximity labelling system to identify interaction partners of Nek2A. Image created with BioRender. (**B**) Cellular localization of TurboID-Nek2A verified by IF staining. (**C**) Plot showing enriched biotinylated proteins identified by Mass-Spec. Data was generated by MaxQuant LFQ analysis. Targets selected for further analysis were marked with blue colour, known interaction partners of Nek2A was colored green. (**D**) Cellular Component analysis on identified peptides in both WT and KD Nek2A interactome indicating sub-cellular localizations. FDR: false discovery rate. (**E, F**) Percentage of multipolar metaphases observed in varying conditions for selected targets, NuMa and KIFC1. Experiments were performed using nocodazole and Plk4 CA models in U2OS and endogenously extra centrosome harbouring cell line SU86.86. (**G**) Pan-cancer Analysis of Advanced and Metastatic Tumors data retrieved from cBioPortal demonstrates positive correlation between NEK2 and KIF2C expressions. (**H**) Co-IP assay to verify physical interaction between Nek2A and KIF2C. (**I**) IF staining demonstrating the co-localization of Nek2A and KIF2C in centrosomes and spindle poles. Arrowheads point to spindle poles and centrosomes where KIF2C and Nek2A co-localize. (**J**) Percentage of multipolar spindles observed when KIF2C is silenced. Experiments were performed as two biological repeats with at least 200 metaphases with CA scored per experiment. . Error bars show standard deviations. Statistical significance was shown as * : p<0.05, **: p<0.01, ***: p<0.001, **** : p<0.0001. KD: kinase-dead, WT: wild type, OX: overexpression

Finding KIFC1 and NuMA, known centrosome clustering factors, led us to explore their collaboration with Nek2A in this process. We used siRNA to silence NuMA (**Supp. Fig. 5C**) and consistent with prior reports [20], NuMA knockdown led to a marked reduction in the formation of MPS. Nevertheless, Nek2A overexpression was capable of inducing MPS without NuMA across three different experimental conditions, implying that Nek2A may govern a mechanism distinct from that of NuMA (**Fig. 5E**). Although NuMA silencing didn’t impede Nek2A’s overexpression impact, there was a competitive interaction during the formation of MPS in metaphase, indicating they regulate centrosome clustering independently and antagonistically. Reversely, silencing Nek2A in NuMA-overexpressing cells also reduced centrosome clustering, further supporting their independent roles (**Supp. Fig. 5D-E**). Next, we investigated how Nek2A interacts with KIFC1, another key centrosome clustering regulator. As reported earlier, silencing KIFC1 impaired centrosome clustering and increased multipolarity [21,22]. Nek2A overexpression in KIFC1-silenced cells raised the percentage of multipolar metaphases, indicating that KIFC1 and Nek2A operate via distinct molecular pathways to orchestrate centrosome clustering (**Fig. 5F**). Supportingly, RNAi against Nek2A partially restored centrosome clustering in KIFC1-deficient cells (**Supp. Fig. 5F-G**).

Analyzing proximity labeling data promoted us to study KIF2C’s role, a kinesin family member. Intriguingly, analysis of cancer patient datasets revealed a strong correlation between the expressions of Nek2A and KIF2C (Spearman: 0.75, Pearson: 0.79, R^2^: 0.62) (**Fig. 5G**), suggesting a possible interaction between these proteins. Confirmatively, ectopically expressed FLAG-tagged Nek2A in U2OS cells showed clear co-immunoprecipitation with KIF2C (**Fig. 5H**), and an anti-KIF2C antibody pulldown also captured Nek2A, verifying their physical interaction. Furthermore, immunofluorescent staining corroborated the colocalisation of Nek2A and KIF2C at spindle poles **(Fig. 5I)**. Overall, while the exact nature of this interaction remains unclear, it’s evident that they associate closely and interact with each other, particularly in the vicinity of the centrosome.

As a novel Nek2A interactor, we assessed the impact of KIF2C knockdown in MPS formation in our CA models **(Fig. 5J)**. KIF2C suppression significantly reduced multipolar metaphases in both models and cell lines. Notably, KIF2C depletion counteracted the effect of Nek2A overexpression, leading to reduced centrosome unclustering. Our data suggests that the interaction between KIF2C and Nek2A is essential to regulate the clustering of extra centrosomes during metaphase.

Further experiments were designed to determine if KIF2C ablation could impede the activity of Nek2A overexpression. We engineered U2OS cells with Nek2A overexpression, KIF2C shRNA expression and a combination of both in addition to Plk4 overexpression to induce CA **(Supp. Fig. 6A)**. In line with our earlier approach, competition experiments showed that suppressing KIF2C enhanced cell viability and proliferation, as evidenced by a marked reduction in multipolar metaphases **(Fig. 6A and Supp. Fig. 6B)**. Interestingly, overexpressing Nek2A did not result in multipolar metaphases, and consequently, cell death was avoided in cells treated with KIF2C shRNA. These observations were substantiated by cell viability assays **(Fig. 6B)** and Annexin-V staining **(Fig. 6C and Supp. Fig. 6C)**. Collectively, our data supports that KIF2C interaction is crucial for Nek2A’s role in promoting the unclustering of extra centrosomes, summarized and presented in **(Fig. 6D)**.

**Figure 6:**
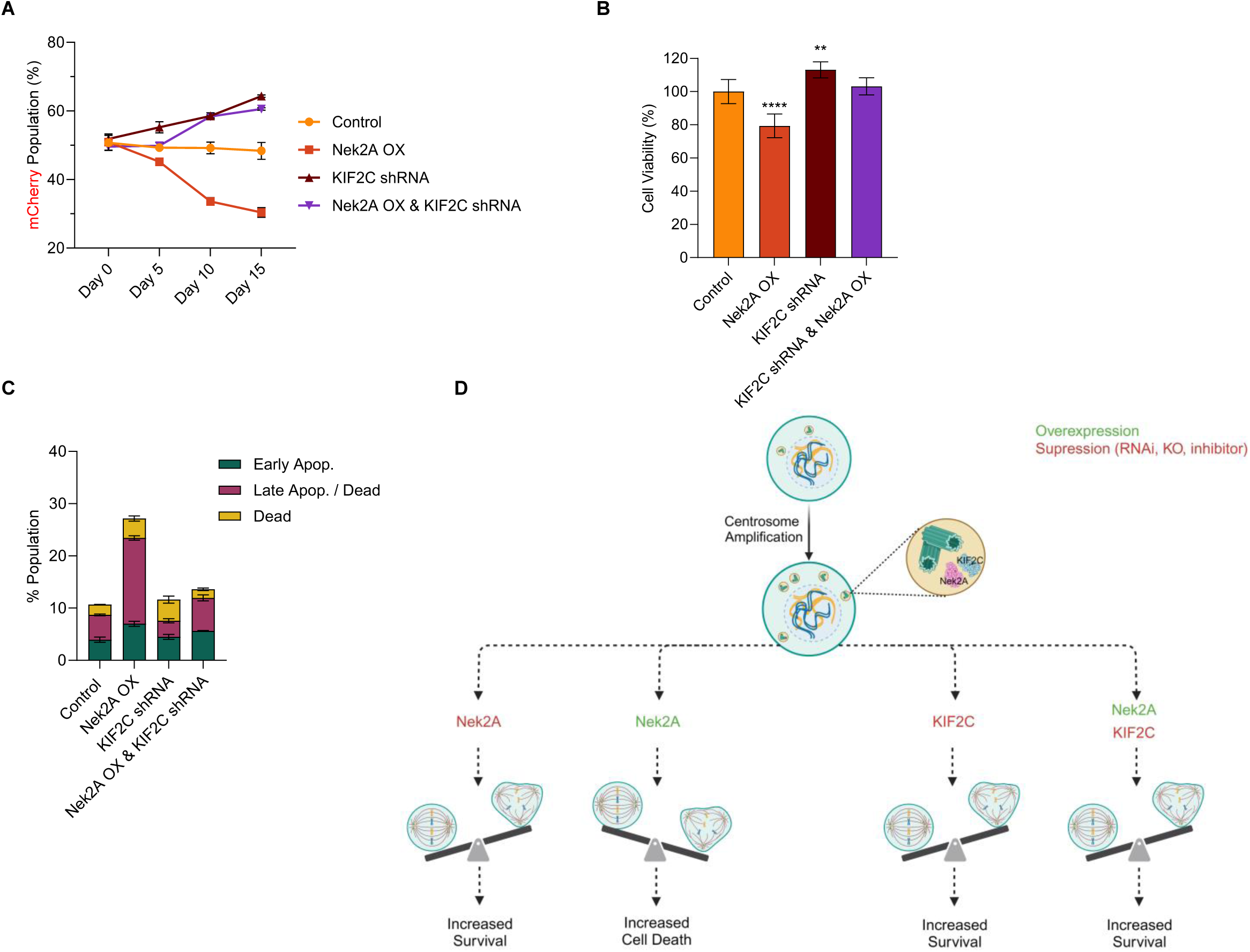
KIF2C is required for Nek2A to exert centrosomal unclustering activity. (**A**) Competition experiment result displays that suppression of KIF2C promotes cell survival in cells with supernumerary centrosomes. (**B**) WST-1 cell viability assay shows that suppression of KIF2C expression increases cell survival and attenuates the effect of Nek2A overexpression on survival of cells harbouring extra centrosomes induced by Plk4 overexpression. (**C**) Annexin-V staining shows that suppression of KIF2C reverts apoptotic phenotype in Nek2A overexpressing cells harbouring Plk4-induced extra centrosomes. (**D**) Schematic representation of the data demonstrating that Nek2A and KIF2C interaction in centrosomes and spindle poles regulate centrosome clustering and affect cancer cell survival. Image created with BioRender. Experiments were performed as two biological repeats. Error bars show standard deviations. Statistical significance was shown as * : p<0.05, **: p<0.01, ***: p<0.001, **** : p<0.0001. OX: overexpression

To link our research to clinical applications, we investigated how high Nek2A levels and CA affect patient survival. Due to the lack of existing data on CA levels and gene expression in patient tumors, we initially analyzed the expression of five centrosomal genes (PLK4, CCNE1 (cyclin E), CETN2 (centrin-2), TUBG1 (γ-tubulin), and PCNT2 (pericentrin)), known as biomarkers for CA [23]. We modified and utilized a previously reported centrosome amplification index (CAI) as outlined in our methods section [23]. This involved using these genes’ expression levels to determine the effect of elevated Nek2A expression on the survival of patients with high CAI. Our results revealed that patients with head and neck squamous cell carcinoma (HNSC), who had both high CAI and high Nek2A levels experienced notably better survival compared to those with high CAI but low Nek2A levels (**Fig. 7A**). Conversely, patients with low CAI experienced poorer outcomes when Nek2A levels were higher, aligning with its established phenotypes [24-27] (**Fig. 7A, left panel**). Next, we hypothesized that chemotherapy with microtubule inhibitors, similar to our nocodazole-induced amplification, might increase centrosome numbers. We then studied taxane response in patients with various Nek2A levels, finding those with higher levels responded better in several cancers (**Fig. 7B and Supp. Table 1**). In these cohorts, we also analyzed the expression levels of CA biomarkers (PLK4, TUBG1, and CCNE1) and found higher expressions in patients who responded well to taxane treatment (**Supp. Fig. 7**). Thus, although various elements affect the response to chemotherapy, the levels of Nek2A stand out as one of the predictors, likely as a result of its unclustering activity. Hence, although Nek2A seems like a promising target, its application must be carefully tailored based on the wider cellular environment, especially considering the level of centrosome amplification.

**Figure 7:**
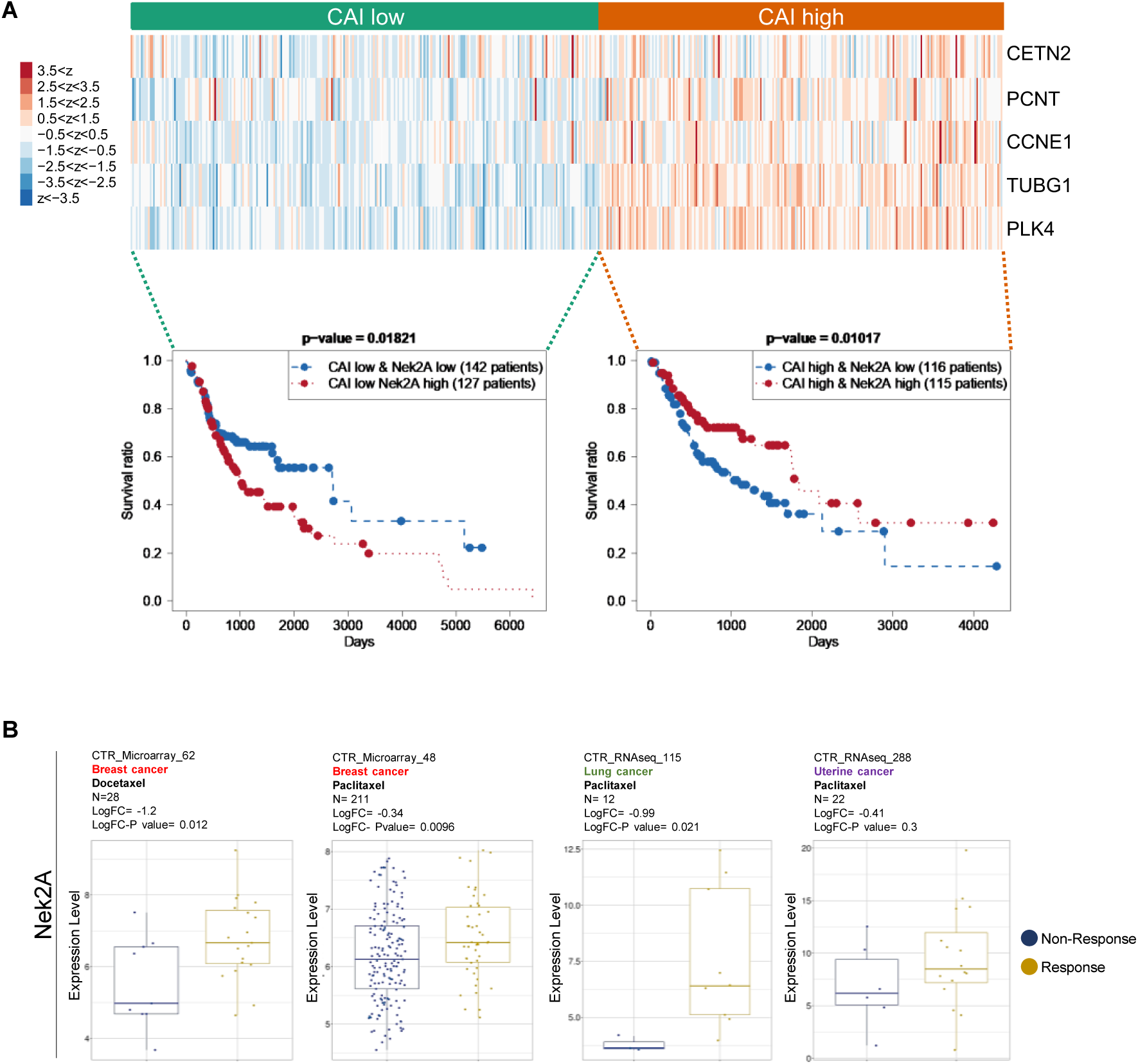
Patient-derived clinical transcriptome data provides evidence that higher Nek2A expression improves survival and taxane response in patients with CA signatures. (**A**) TCGA data analysis indicates that high Nek2A expression significantly increases the survival rate of HNSC patients with high centrosome amplification index (CAI). (**B**) In cohorts of cancer patients exhibiting positive response to taxane treatment, relatively higher expression levels of Nek2A is observed compared to the non-responsive patients. Data was obtained from Cancer Treatment Response Gene Signature Data Base (CTR-DB).

## DISCUSSION

Overexpression of Nek2A has been long recognized as a facilitator of tumor growth, migration, and drug resistance, marking it as a prognostic indicator and a potential target in anticancer therapy [27-31]. In our study, we explored the role of Nek2A in cells possessing extra centrosomes. We discovered that overexpression of Nek2A not only unclustered these supernumerary centrosomes, but also decreased cell viability and increased apoptosis, likely due to the induction of multipolar divisions. Multipolar spindle formation and subsequent cell death occurred selectively in cells with extra centrosomes, while the viability of cells without centrosome amplification remained unaffected. These findings suggest that the impact of Nek2A targeting can vary depending on the genetic and cellular context of the cancer cells, a consideration that has been largely neglected until now. Supporting this argument, HNSC patients with higher levels of CA and Nek2A demonstrated improved survival rates. Moreover, breast cancer patients with increased Nek2A expression had a more positive response to taxane treatment. We acknowledge that while this improvement could be due to centrosome amplification after taxane treatment, akin to our experimental results with nocodazole, other factors could also play a role.

Investigating Nek2A’s role in centrosomal unclustering, we initially focused on potential centrosomal targets like C-Nap1, Rootletin, and Gas2L1. Consistent with literature, we found that the loss of C-Nap1 disrupts centrosome organization, increases centrosome distance, and leads to multipolar divisions [18,19]. This effect is further amplified by KIFC1 inhibition in C-Nap1 knockout cells, indicating an independent role for C-Nap1 in centrosome clustering. Similarly, overexpressing Nek2A in these cells enhances multipolar metaphases, pointing to a synergistic effect with C-Nap1 in this process. Furthermore, Nek2A overexpression significantly increases multipolarity in cells lacking Rootletin, underscoring its influence beyond C-Nap1. Lastly, our findings suggest that GAS2L1, despite being a recent target of Nek2A [15], does not significantly impact centrosome clustering. This suggests that the unclustering effect of Nek2A overexpression is independent of the specific centrosomal targets during the G2/M transition.

Shifting focus to Nek2A’s broader interactions, we explored its phosphorylation of mitotic proteins such as Hec1. Using INH154 to inhibit Nek2A-Hec1 interaction revealed that Hec1 is not essential for Nek2A-induced multipolar metaphases. Additionally, our study found that Nek2A’s interaction with TRF1, though influential in cytokinesis in cells with centrosome amplification [17,32], is not crucial for centrosome clustering. This comprehensive analysis leads us to propose that the unclustering effect of Nek2A overexpression involves novel targets, expanding the scope of Nek2A’s impact in cellular processes.

In our study, we utilized TurboID for efficient proximity labeling to explore Nek2A’s interaction partners in centrosome clustering. Analysis identified 20 significantly enriched and previously known (BioGrid & IntAct databases) Nek2A interaction partners, including Nek2, with similar enrichment in both WT and KD pull-downs, suggesting that kinase function does not greatly change Nek2’s interactome. Despite finding clustering regulators NuMA and KIFC1 within Nek2A-TurboID’s labeling radius, our results indicate that Nek2A’s centrosome unclustering activity is independent of these proteins. Notably, co-immunoprecipitation experiments showed that Nek2A and NuMA do not physically interact, each regulating centrosome clustering independently.

Moreover, KIFC1, essential for centrosome clustering in cancer cells [22,33], is found to be functionally independent from Nek2A. Our experiments revealed that Nek2A overexpression increases multipolar spindles formation even without KIFC1, and their co-suppression partially restores centrosome clustering. This suggests that Nek2A and KIFC1 independently control centrosome clustering, acting as competitors in this process, a finding supported by the positive correlation between their expressions. This reveals a complex balancing mechanism where overexpression of both proteins induces contrasting effects on centrosome clustering.

KIF2C, a member of the kinesin-13 family known for regulating microtubule dynamics [34], has not been extensively studied in the context of centrosome clustering in human cancer. Recent findings indicate that both knockdown and overexpression of KIF2C in HeLa cells increase multipolar metaphases, suggesting its activity is tightly regulated during mitosis [35]. Key regulatory mechanisms include phosphorylation by Aurora B, which inhibits KIF2C while guiding its centromere localization, and Aurora A [36-38]. Additionally, Cdk1 phosphorylation releases KIF2C from centrosomes and modulates its depolymerizing activity, while Plk1 phosphorylates both KIF2C and Nek2A, the latter promoting centrosome splitting [39-42].

Our study reveals a novel interaction between Nek2A and KIF2C in cancer cells with centrosome amplification, suggesting a coordinated regulation to maintain centrosome clustering. This interaction, validated through proximity labeling, co-staining, and co-immunoprecipitation, highlights the intricate relationship between mitotic kinases and kinesins in modulating microtubule dynamics and centrosome clustering during mitosis.

In this study, we have discovered a critical interaction between KIF2C and Nek2A that regulates centrosome clustering in cancer cells, marking the first identification of such a relationship. This interaction is most likely a kinase-substrate type, as evidenced by our findings where overexpression of a kinase-dead Nek2A mutant acted dominantly negative, indicating the necessity of kinase activity for phenotypical outcomes. Further research is needed to elucidate the specifics of the KIF2C-Nek2A interaction.

## CONCLUSION

In conclusion, this study illuminates the crucial role of Nek2A in controlling centrosome clustering in cancer cells, particularly those with extra centrosomes. Overexpression of Nek2A represents a disadvantage for cancer cells with extra centrosomes, shown under both *in vitro and in vivo* conditions. Silencing KIF2C rescues the cells from the detrimental effect of Nek2A. This study highlights the potential of targeting Nek2A and its interactors, like KIF2C, for novel cancer therapies aimed at managing centrosome-related genomic instability. Further exploration of these molecular interactions promises to enhance our understanding of centrosome clustering regulation and its significance in cancer biology, opening new avenues for effective cancer treatment strategies.

## MATERIAL AND METHODS

### Cell Culture

N1E115 (ATCC, CRL-2263), MDA-MB-231 (ATCC, HTB-26), U2OS (ATCC, HTB-96), SU86.86 (ATCC, CRL-1837), MIA PaCa-2 (ATCC, CRL-1420), Panc1 (ATCC, CRL-1469), HEK293T (ATCC, CRL-3216) were maintained in DMEM (Gibco) supplemented with 10% FBS (Gibco) and 1% Pen/Strep (Gibco) at 37°C and 5% CO_2_ incubator.

### Plasmids

Nek2A (clone ID: 38963) in pJP1563 and NuMA1 (clone ID: 871325) in pLenti6.3/V5-DEST were purchased from DNASU. LentiCRISPR-v2 (52961), pCW57-RFP-P2A-MCS (78933), pCW57-MCS1-P2A-MCS2 (80922), pCDH-EF1-FHC (64874), Flag-TurboID (124646), Tet-pLKO-neo (21916), pcDNA3-Plk4 (41165), pcDNA5-STIL (80266), PGK-H2B-mCherry (21217) and PGK-H2B-eGFP (21210) were purchased from Addgene.

Kinase-dead mutant (K37R) of Nek2A was derived from Nek2A in pJP1563 using Q5 Site-Directed Mutagenesis Kit (NEB) following the instructions of manufacturer. Oligos used for the SDM reaction are given in (**Supp. Table 2**). Single guide RNA (sgRNA) oligos (**Supp. Table 3**) for CRISPR/Cas9 knockout of Nek2A, C-Nap1 and Rootletin were cloned into LentiCRISPR-v2 as previously described [43]. Dox-inducible overexpressions of Nek2A and PLK4 were achieved by subcloning to pCW57-RFP-P2A-MCS and pCW57-MCS1-P2A-MCS2 respectively. Nek2A-WT and Nek2A-KD(K37R) cDNAs were subcloned to Flag-TurboID. To perform Co-IP using anti-FLAG antibody, Nek2A cDNA was subcloned into pCDH-EF1-FHC. Oligos for dox-inducible shRNA expression targeting KIF2C were cloned into Tet-pLKO-neo as previously described [44] and provided in (**Supp. Table 4**).

### Transfection and Viral Transduction

Nek2A, Gas2L1, TRF1, NuMA, KIFC1 and KIF2C were silenced by siRNA transfections. List of the siRNAs is provided in (**Supp. Table 5**). Briefly, 2×10^5^ cells were seeded in 6 well plates, 100 pmol siRNA, and 7,5 µl of Lipofectamine 3000 (Thermo) were added to culture media according to the manufacturer’s protocol. Knock-down efficiencies were analysed by either Western Blot or RT-qPCR or both.

Overexpressions (Nek2A, PLK4, NuMA), CRISPR/Cas9-mediated knockouts and shRNAs were delivered to target cells via viral particles. To generate viral particles, 2×10^6^ HEK293T cells were seeded per 10cm petri dish. 2500 ng transfer vector, 2250 ng packaging vector (psPAX2 for lentivirus, pUMVC for retrovirus) and 250 ng envelope vector (pCMV-VSV-G) are mixed with 20 µL Fugene 6 (Roche, USA) diluted in OptiMEM. Cells were transfected with the mixture prepared. Culture medium was collected 48-and 72-hours post-transfection and 100X concentrated by 50% (w/v) PEG 8000 (P2139, Sigma). Infections were performed using 10 µL virus and 8 µg/ml protamine sulphate in 2 ml culture medium.

### Centrosome amplification using microtubule inhibitor

2×10^5^ cells were seeded on 15×15 mm coverslips. Nocodazole (100 ng/ml) treatment was done for 16 hours to achieve prometaphase arrest and mitotic slip, resulting in duplicated centrosomes. Culture media was replenished without nocodazole, and cells were incubated 24 hours to allow cells to recover and re-enter the cell cycle with amplified centrosomes.

### Immunofluorescence staining

Cells grown on coverslips were fixed with ice-cold methanol for 15 minutes, washed 3 times with DPBS-T, and blocked with 5% (w/v) BSA for 30 minutes. Samples were incubated with primary antibodies for γ-Tubulin (SIGMA, T5192), α-Tubulin (Abcam, 7291), Nek2 (BD Biosciences, 610593), FLAG (SIGMA, F1804), KIF2C (Santa Cruz, sc-81305) (1:500 dilution in 1% BSA in PBS) at 4°C overnight, followed by incubation at RT for 2 hours with secondary antibodies (1:1000 dilution in 1% BSA in PBS) (Alexa flour 488 and 594, Thermo). Cell nuclei were labelled by DAPI containing mounting medium (Vectashield).

### RT-qPCR

Total RNA is isolated using NucleoSpin RNAII kit (Macherey-Nagel) following the manufacturer’s instructions. 1000 ng total RNA is reverse transcribed using M-MLV Reverse Transcriptase (Invitrogen) cDNA synthesis kit following the manufacturer’s protocol. RT-qPCR is performed using The LightCycler 480 SYBR Green I (Roche), and the reaction was run at LightCycler 480 Instrument II (Roche). Samples were normalised to GAPDH expression. The PCR products are subjected to a melting curve analysis. Primers used in this study are provided in (**Supp. Table 6**).

### Western Blot

Cell pellets were lysed in RIPA buffer (50 mM Tris, pH 7.4, 500 mM NaCI, 0.4% SDS, 5 mM EDTA, 1 mM DTT, 2% Triton X-100, with protease and phosphatase inhibitors). Samples are run on SDS-PAGE for separation and then transferred to PVDF membranes. Membranes are blocked with 5% non-fat dry milk in 1X TBS-T and then incubated overnight at 4°C with primary antibodies with recommended or optimised dilutions. Membranes were washed 3 times with TBS-T and incubated with corresponding secondary antibodies at RT for 2 hours. Blots are visualised using the Licor Odyssey FC imaging system. Beta-actin was used as the loading control.

### Metaphase Scoring

Cells were stained for γ-Tubulin and DAPI and a minimum of 300 metaphases with CA/experiment were scored and each experiment was repeated at least twice. CA was measured based on centrosome number per cell at interphase. Cells bearing >2 centrosomes were marked as CA.

### Cell Viability

Cell viability was determined with WST-1 assay: 3,000 cells/well were seeded in 96-well plates, adhered overnight (16 hours), followed by a 30-minute incubation with WST-1 reagent (Roche) and absorbance read at 440 nm using a microplate reader (Synergy H1 Reader, Biotek, USA).

### Cell Cycle and Apoptosis Assays

Cell cycle analysis was performed using the Muse Cell Cycle Assay Kit (Millipore). The Annexin V staining procedure was conducted using the Muse® Annexin V & Dead Cell Kit from Luminex (MCH100105), following the provided manufacturer’s guidelines. Flow cytometry analysis was conducted using the Muse Cell Analyzer, and the data were processed using Muse analysis software.

### Dual-Color Competition Assays

The long-term effects on cell survival and proliferation were assessed using dual-color competition assays *in vitro* and *in vivo*. Cells were tagged with either mCherry or eGFP and mixed in a 1:1 ratio. For *in vitro* assays, 5 x 10⁴ cells were seeded in 6-well plates and passaged 1:5 at confluence. Plates were imaged at day 0, 5, 10, and 15 using an Agilent BioTek Cytation 5 imaging platform. mCherry-positive cells were quantified using Cytation 5 software. For *in vivo* assays, 100 µl of a 1:1 mixture of 2 x 10⁷ cells/ml in Matrigel was injected subcutaneously into SCID mice. After 8 weeks, 6 tumors were collected, fixed, dehydrated, cleared, and paraffin-embedded. 2 µm sections were obtained using microtome (Leica RM2245) picked randomly to represent different parts of the tumor.

### TurboID Proximity Labelling

Cell pellets were obtained by centrifugation following biotinylation and PBS washes. Protein lysates were prepared by incubating cell pellets with protease inhibitor in lysis buffer. Protein concentrations were determined using the BCA method, and proteins were subjected to overnight incubation with Streptavidin beads at 4°C. Beads were washed twice to remove unbound proteins, and bound proteins were subsequently subjected to trypsin digestion. Following digestion, formic acid was added to the samples to achieve a final concentration of 5%. The resulting supernatants were collected and analyzed using a Thermo Scientific Q Exactive HF Hybrid Quadrupole-Orbitrap Mass Spectrometer. Peptide identification and quantification were performed using Thermo Fisher Scientific Proteome Discoverer and MaxQuant software. Known contaminant proteins, such as keratin, were excluded from the analysis, and proteins identified with at least two unique peptides were considered significant in terms of abundance. The mass spectrometry proteomics data have been deposited to the ProteomeXchange Consortium via the PRIDE [45] partner repository with the dataset identifier PXD046867.

### Co-Immunoprecipitation (Co-IP)

Cells were harvested via trypsinization, crosslinked with 0.1% paraformaldehyde (PFA) for 7 minutes, and lysed directly in ice-cold immunoprecipitation (IP) buffer supplemented with protease inhibitor, phosphatase inhibitor, and PMSF. Protein concentration was determined using a BCA assay. Protein G magnetic beads were pre-cleared and incubated with protein samples and specific antibodies or IgG control for 2 hours at 4°C. Following overnight incubation with protein G magnetic beads, beads were washed three times with IP buffer, and proteins were eluted by denaturation with 1X Laemmli buffer containing 50 mM DTT at 95°C for 15 minutes. Western blotting was employed for subsequent analysis.

### Microscopy

Leica DMI8 SP8 microscope and LASX software was used for confocal imaging and image processing. Metaphase were scored using Carl Zeiss Axio Imager M1. Competition assays were performed by using BioTek Cytation 5 Cell Imaging Multimode Reader.

### Patient Data Analysis

Gene expression data for tumors were processed using The Cancer Genome Atlas (TCGA) consortium’s RNA-Seq pipeline (https://portal.gdc.cancer.gov). We downloaded HTSeq-FPKM files for all primary tumors from the latest data release (Data Release 38–August 31, 2023), excluding metastatic tumors due to their distinct biology. Patient survival data were extracted from clinical annotation files. Our survival analysis included only patients with both survival data and gene expression profiles. We log_2_-transformed and *z*-normalized the gene expression profiles within each cohort, generating heat maps from these normalized values. For gene set analyses, we used *k*-means clustering (*k* = 2) on the normalized data, comparing Kaplan-Meier survival curves for the two groups via a log-rank test. We further divided each group based on Nek2A expression to examine its impact on survival.

CTR-DB (Cancer Treatment Response gene signature DataBase) [46] was used to process patient transcriptome data along with taxane response and expression levels of Nek2A, KIF2C and centrosome amplification biomarkers PLK4, TUBG1 and CCNE1.

### Statistical Analysis

All experiments were conducted as biological repeats, and statistical analysis was performed using GraphPad Prism version 9.0. The student’s t-test was employed to compare two groups, while the two-way ANOVA was used to compare more than two groups for parametric variables. Significance levels were denoted as follows: * for p<0.05, ** for p<0.01, and *** for p<0.001.

## ACKNOWLEDGEMENTS

The authors gratefully acknowledge the use of the services and facilities of the Koç University Research Center for Translational Medicine (KUTTAM), funded by the Presidency of Turkey, Presidency of Strategy and Budget. The authors express sincere gratitude to William S. Saunders and Nicholas Quintyne for their intellectual contributions. Illustrations in the figures were created by using BioRender.com and licensed (3P9YOS, XR263PAA8S, XL264EHWDK, JS263OX89L, TX263PBZZZ, LN263P92LT) for publication purposes.

## REFERENCES

1. Song, S.; Jung, S.; Kwon, M. Expanding roles of centrosome abnormalities in cancers. BMB Rep 2023, 56, 216–224, doi:10.5483/BMBRep.2023-0025.

2. Kalkan, B.M.; Ozcan, S.C.; Quintyne, N.J.; Reed, S.L.; Acilan, C. Keep Calm and Carry on with Extra Centrosomes. Cancers 2022, 14, 442.

3. Godinho, S.A.; Pellman, D. Causes and consequences of centrosome abnormalities in cancer. Philos Trans R Soc Lond B Biol Sci 2014, 369, doi:10.1098/rstb.2013.0467.

4. Fry, A.M.; Meraldi, P.; Nigg, E.A. A centrosomal function for the human Nek2 protein kinase, a member of the NIMA family of cell cycle regulators. The EMBO journal 1998, 17, 470–481.

5. Fry, A.M. The Nek2 protein kinase: a novel regulator of centrosome structure. Oncogene 2002, 21, 6184–6194, doi:10.1038/sj.onc.1205711.

6. O’regan, L.; Blot, J.; Fry, A.M. Mitotic regulation by NIMA-related kinases. Cell division 2007, 2, 1–12.

7. Fry, A.M.; Mayor, T.; Meraldi, P.; Stierhof, Y.-D.; Tanaka, K.; Nigg, E.A. C-Nap1, a novel centrosomal coiled-coil protein and candidate substrate of the cell cycle–regulated protein kinase Nek2. The Journal of cell biology 1998, 141, 1563–1574.

8. Bahe, S.; Stierhof, Y.-D.; Wilkinson, C.J.; Leiss, F.; Nigg, E.A. Rootletin forms centriole-associated filaments and functions in centrosome cohesion. The Journal of cell biology 2005, 171, 27–33.

9. Ring, D.; Hubble, R.; Kirschner, M. Mitosis in a cell with multiple centrioles. The Journal of Cell Biology 1982, 94, 549–556.

10. Mittal, K.; Ogden, A.; Reid, M.D.; Rida, P.C.G.; Varambally, S.; Aneja, R. Amplified centrosomes may underlie aggressive disease course in pancreatic ductal adenocarcinoma. Cell Cycle 2015, 14, 2798–2809, doi:10.1080/15384101.2015.1068478.

11. Faragher, A.J.; Fry, A.M. Nek2A kinase stimulates centrosome disjunction and is required for formation of bipolar mitotic spindles. Mol Biol Cell 2003, 14, 2876–2889, doi:10.1091/mbc.e03-02-0108.

12. Mayor, T.; Hacker, U.; Stierhof, Y.D.; Nigg, E.A. The mechanism regulating the dissociation of the centrosomal protein C-Nap1 from mitotic spindle poles. J Cell Sci 2002, 115, 3275–3284, doi:10.1242/jcs.115.16.3275.

13. Yang, J.; Adamian, M.; Li, T. Rootletin interacts with C-Nap1 and may function as a physical linker between the pair of centrioles/basal bodies in cells. Molecular biology of the cell 2006, 17, 1033–1040.

14. Au, F.K.; Jia, Y.; Jiang, K.; Grigoriev, I.; Hau, B.K.; Shen, Y.; Du, S.; Akhmanova, A.; Qi, R.Z. GAS2L1 is a centriole-associated protein required for centrosome dynamics and disjunction. Developmental cell 2017, 40, 81–94.

15. Au, F.K.C.; Hau, B.K.T.; Qi, R.Z. Nek2-mediated GAS2L1 phosphorylation and centrosome-linker disassembly induce centrosome disjunction. Journal of Cell Biology 2020, 219, doi:10.1083/jcb.201909094.

16. Chen, Y.; Riley, D.J.; Zheng, L.; Chen, P.-L.; Lee, W.-H. Phosphorylation of the mitotic regulator protein Hec1 by Nek2 kinase is essential for faithful chromosome segregation. Journal of Biological Chemistry 2002, 277, 49408–49416.

17. Prime, G.; Markie, D. The telomere repeat binding protein Trf1 interacts with the spindle checkpoint protein Mad1 and Nek2 mitotic kinase. Cell Cycle 2005, 4, 121–124.

18. Panic, M.; Hata, S.; Neuner, A.; Schiebel, E. The centrosomal linker and microtubules provide dual levels of spatial coordination of centrosomes. PLoS Genet 2015, 11, e1005243, doi:10.1371/journal.pgen.1005243.

19. Theile, L.; Li, X.; Dang, H.; Mersch, D.; Anders, S.; Schiebel, E. Centrosome linker diversity and its function in centrosome clustering and mitotic spindle formation. Embo j 2023, e109738, doi:10.15252/embj.2021109738.

20. Quintyne, N.J.; Reing, J.E.; Hoffelder, D.R.; Gollin, S.M.; Saunders, W.S. Spindle multipolarity is prevented by centrosomal clustering. Science 2005, 307, 127–129, doi:10.1126/science.1104905.

21. Leber, B.; Maier, B.; Fuchs, F.; Chi, J.; Riffel, P.; Anderhub, S.; Wagner, L.; Ho, A.D.; Salisbury, J.L.; Boutros, M. Proteins required for centrosome clustering in cancer cells. Science translational medicine 2010, 2, 33ra38–33ra38.

22. Kwon, M.; Godinho, S.A.; Chandhok, N.S.; Ganem, N.J.; Azioune, A.; Thery, M.; Pellman, D. Mechanisms to suppress multipolar divisions in cancer cells with extra centrosomes. Genes Dev 2008, 22, 2189–2203, doi:10.1101/gad.1700908.

23. Pannu, V.; Mittal, K.; Cantuaria, G.; Reid, M.D.; Li, X.; Donthamsetty, S.; McBride, M.; Klimov, S.; Osan, R.; Gupta, M.V. Rampant centrosome amplification underlies more aggressive disease course of triple negative breast cancers. Oncotarget 2015, 6, 10487.

24. Xu, H.; Zeng, L.; Guan, Y.; Feng, X.; Zhu, Y.; Lu, Y.; Shi, C.; Chen, S.; Xia, J.; Guo, J.;, et al. High NEK2 confers to poor prognosis and contributes to cisplatin-based chemotherapy resistance in nasopharyngeal carcinoma. J Cell Biochem 2019, 120, 3547–3558, doi:10.1002/jcb.27632.

25. Wang, C.; Huang, Y.; Ma, X.; Wang, B.; Zhang, X. Overexpression of NEK2 is correlated with poor prognosis in human clear cell renal cell carcinoma. Int J Immunopathol Pharmacol 2021, 35, 20587384211065893, doi:10.1177/20587384211065893.

26. Zeng, Y.-R.; Han, Z.-D.; Wang, C.; Cai, C.; Huang, Y.-Q.; Luo, H.-W.; Liu, Z.-Z.; Zhuo, Y.-J.; Dai, Q.-S.; Zhao, H.-B. Overexpression of NIMA-related kinase 2 is associated with progression and poor prognosis of prostate cancer. BMC urology 2015, 15, 1–8.

27. Kokuryo, T.; Yokoyama, Y.; Yamaguchi, J.; Tsunoda, N.; Ebata, T.; Nagino, M. NEK2 is an effective target for cancer therapy with potential to induce regression of multiple human malignancies. Anticancer research 2019, 39, 2251–2258.

28. Li, Y.; Chen, L.; Feng, L.; Zhu, M.; Shen, Q.; Fang, Y.; Liu, X.; Zhang, X. NEK2 promotes proliferation, migration and tumor growth of gastric cancer cells via regulating KDM5B/H3K4me3. Am J Cancer Res 2019, 9, 2364–2378.

29. Bai, R.; Yuan, C.; Sun, W.; Zhang, J.; Luo, Y.; Gao, Y.; Li, Y.; Gong, Y.; Xie, C. NEK2 plays an active role in Tumorigenesis and Tumor Microenvironment in Non-Small Cell Lung Cancer. Int J Biol Sci 2021, 17, 1995–2008, doi:10.7150/ijbs.59019.

30. Zhou, W.; Yang, Y.; Xia, J.; Wang, H.; Salama, M.E.; Xiong, W.; Xu, H.; Shetty, S.; Chen, T.; Zeng, Z. NEK2 induces drug resistance mainly through activation of efflux drug pumps and is associated with poor prognosis in myeloma and other cancers. Cancer cell 2013, 23, 48–62.

31. Cusan, M.; Wang, L. NEK2, a promising target in TP53 mutant cancer. Blood Science 2022, 04, 97–98, doi:doi:10.1097/BS9.0000000000000106.

32. Ohishi, T.; Muramatsu, Y.; Yoshida, H.; Seimiya, H. TRF1 ensures the centromeric function of Aurora-B and proper chromosome segregation. Molecular and cellular biology 2014, 34, 2464–2478.

33. Watts, C.A.; Richards, F.M.; Bender, A.; Bond, P.J.; Korb, O.; Kern, O.; Riddick, M.; Owen, P.; Myers, R.M.; Raff, J. Design, synthesis, and biological evaluation of an allosteric inhibitor of HSET that targets cancer cells with supernumerary centrosomes. Chemistry & biology 2013, 20, 1399–1410.

34. Ritter, A.; Kreis, N.-N.; Louwen, F.; Wordeman, L.; Yuan, J. Molecular insight into the regulation and function of MCAK. Critical Reviews in Biochemistry and Molecular Biology 2016, 51, 228–245.

35. Moon, H.H.; Kreis, N.N.; Friemel, A.; Roth, S.; Schulte, D.; Solbach, C.; Louwen, F.; Yuan, J.; Ritter, A. Mitotic Centromere-Associated Kinesin (MCAK/KIF2C) Regulates Cell Migration and Invasion by Modulating Microtubule Dynamics and Focal Adhesion Turnover. Cancers (Basel) 2021, 13, doi:10.3390/cancers13225673.

36. Andrews, P.D.; Ovechkina, Y.; Morrice, N.; Wagenbach, M.; Duncan, K.; Wordeman, L.; Swedlow, J.R. Aurora B regulates MCAK at the mitotic centromere. Developmental cell 2004, 6, 253–268.

37. Lan, W.; Zhang, X.; Kline-Smith, S.L.; Rosasco, S.E.; Barrett-Wilt, G.A.; Shabanowitz, J.; Hunt, D.F.; Walczak, C.E.; Stukenberg, P.T. Aurora B phosphorylates centromeric MCAK and regulates its localization and microtubule depolymerization activity. Current Biology 2004, 14, 273–286.

38. Zhang, X.; Ems-McClung, S.C.; Walczak, C.E. Aurora A phosphorylates MCAK to control ran-dependent spindle bipolarity. Molecular biology of the cell 2008, 19, 2752–2765.

39. Sanhaji, M.; Friel, C.T.; Kreis, N.-N.; Krämer, A.; Martin, C.; Howard, J.; Strebhardt, K.; Yuan, J. Functional and spatial regulation of mitotic centromere-associated kinesin by cyclin-dependent kinase 1. Molecular and cellular biology 2010, 30, 2594–2607.

40. Zhang, W.; Fletcher, L.; Muschel, R.J. The role of Polo-like kinase 1 in the inhibition of centrosome separation after ionizing radiation. Journal of Biological Chemistry 2005, 280, 42994–42999.

41. Zhang, L.; Shao, H.; Huang, Y.; Yan, F.; Chu, Y.; Hou, H.; Zhu, M.; Fu, C.; Aikhionbare, F.; Fang, G. PLK1 phosphorylates MCAK and promotes its depolymerase activity. J Biol Chem 2010.

42. Mbom, B.C.; Siemers, K.A.; Ostrowski, M.A.; Nelson, W.J.; Barth, A.I. Nek2 phosphorylates and stabilizes β-catenin at mitotic centrosomes downstream of Plk1. Mol Biol Cell 2014, 25, 977–991, doi:10.1091/mbc.E13-06-0349.

43. Sanjana, N.E.; Shalem, O.; Zhang, F. Improved vectors and genome-wide libraries for CRISPR screening. Nat Methods 2014, 11, 783–784, doi:10.1038/nmeth.3047.

44. Wiederschain, D.; Wee, S.; Chen, L.; Loo, A.; Yang, G.; Huang, A.; Chen, Y.; Caponigro, G.; Yao, Y.M.; Lengauer, C.;, et al. Single-vector inducible lentiviral RNAi system for oncology target validation. Cell Cycle 2009, 8, 498–504, doi:10.4161/cc.8.3.7701.

45. Perez-Riverol, Y.; Bai, J.; Bandla, C.; García-Seisdedos, D.; Hewapathirana, S.; Kamatchinathan, S.; Kundu, D.J.; Prakash, A.; Frericks-Zipper, A.; Eisenacher, M.;, et al. The PRIDE database resources in 2022: a hub for mass spectrometry-based proteomics evidences. Nucleic Acids Res 2022, 50, D543–d552, doi:10.1093/nar/gkab1038.

46. Liu, Z.; Liu, J.; Liu, X.; Wang, X.; Xie, Q.; Zhang, X.; Kong, X.; He, M.; Yang, Y.; Deng, X.;, et al. CTR-DB, an omnibus for patient-derived gene expression signatures correlated with cancer drug response. Nucleic Acids Res 2022, 50, D1184–d1199, doi:10.1093/nar/gkab860.

