## supplemental tables for "Less is More: Nek2A Unclusters Extra Centrosomes and Induces Cell Death in Cancer Cells via KIF2C Interaction"

**Supplementary Table 1:** Expression levels of Nek2A and KIF2C in different patient datasets. LogFC indicates the comparison of gene expression between responsive and non-responsive individuals against taxane treatment. Data was accessed in Cancer Treatment Response Gene Signature Data Base (CTR-DB)

| | | | | NEK2 | | KIF2C | |
|---|---|---|---|---|---|---|---|
| **CTR-DB ID** | **Cancer Type** | **Drug** | **Sample size** | **LogFC** | **LogFC-P value** | **LogFC** | **LogFC-P value** |
| CTR_Microarray_48 | Breast cancer | Paclitaxel | 211 | -0.34 | 0.0096 | -0.54 | 2.10E-06 |
| CTR_Microarray_75 | Breast cancer | Paclitaxel | 124 | -0.025 | 0.92 | -0.02 | 0.93 |
| CTR_Microarray_106 | Breast cancer | Paclitaxel | 79 | -0.094 | 0.6 | -0.26 | 0.17 |
| CTR_Microarray_104 | Breast cancer | Paclitaxel | 71 | -0.24 | 0.57 | -0.47 | 0.07 |
| CTR_Microarray_34 | Ovarian cancer | Paclitaxel | 64 | 0.0017 | 0.99 | -0.1 | 0.53 |
| CTR_Microarray_28 | Breast cancer | Paclitaxel | 63 | -0.23 | 0.12 | 0.018 | 0.85 |
| CTR_Microarray_40 | Breast cancer | Paclitaxel | 51 | -0.24 | 0.15 | -0.019 | 0.87 |
| CTR_Microarray_72 | Breast cancer | Paclitaxel | 44 | -0.42 | 0.29 | -0.2 | 0.46 |
| CTR_Microarray_1 | Breast cancer | Paclitaxel | 42 | -0.4 | 0.19 | -0.14 | 0.31 |
| CTR_Microarray_55 | Breast cancer | Paclitaxel | 34 | 0.59 | 0.17 | 0.3 | 0.36 |
| CTR_Microarray_47 | Breast cancer | Paclitaxel | 31 | -0.097 | 0.87 | -0.19 | 0.6 |
| CTR_Microarray_41 | Breast cancer | Paclitaxel | 23 | -0.32 | 0.55 | -0.58 | 0.23 |
| CTR_RNAseq_288 | Uterine cancer | Paclitaxel | 22 | -0.41 | 0.3 | -0.17 | 0.54 |
| CTR_Microarray_71 | Ovarian cancer | Paclitaxel | 20 | 0.86 | 0.064 | 0.66 | 0.18 |
| CTR_Microarray_103 | Breast cancer | Paclitaxel | 16 | -0.41 | 0.51 | -0.31 | 0.54 |
| CTR_RNAseq_201 | Uterine cancer | Paclitaxel | 15 | 0.59 | 0.39 | 0.58 | 0.51 |
| CTR_RNAseq_115 | Lung cancer | Paclitaxel | 12 | -0.99 | 0.021 | -0.31 | 0.47 |
| CTR_Microarray_107 | Breast cancer | Docetaxel | 66 | -0.33 | 0.064 | -0.46 | 0.00048 |
| CTR_Microarray_74 | Breast cancer | Docetaxel | 32 | -0.46 | 0.12 | -0.42 | 0.0079 |
| CTR_Microarray_4 | Breast cancer | Docetaxel | 31 | -0.16 | 0.74 | -0.12 | 0.78 |
| CTR_Microarray_96 | Breast cancer | Docetaxel | 29 | -0.078 | 0.78 | -0.37 | 0.066 |
| CTR_Microarray_62 | Breast cancer | Docetaxel | 28 | -1.2 | 0.012 | -0.44 | 0.16 |
| CTR_Microarray_61 | Breast cancer | Docetaxel | 25 | 0.53 | 0.056 | -0.031 | 0.88 |
| CTR_Microarray_98 | Breast cancer | Docetaxel | 21 | -0.12 | 0.84 | 0.17 | 0.71 |
| CTR_RNAseq_272 | Soft tissue tumours | Docetaxel | 16 | -0.2 | 0.77 | -0.35 | 0.62 |
| CTR_Microarray_85 | Breast cancer | Docetaxel | 16 | -0.15 | 0.7 | -0.098 | 0.8 |

**Supplementary Table 2:** Oligo sequences used for site-directed mutagenesis (SDM) reaction to generated Nek2A-(K37R)

| Name | Sequence (5' - 3') |
|---|---|
| NEK2_SDM_K37R-F | TGA TGG CAA GAT ATT AGT TTG GAG AGA ACT TGA CTA TGG CTC |
| NEK2_SDM_K37R-R | GAG CCA TAG TCA AGT TCT CTC CAA ACT AAT ATC TTG CCA TCA |

**Supplementary Table 3:** Oligo sequences for sgRNA cloning into LentiCRISPR-v2

| Name | Sequence (5' - 3') |
|---|---|
| Nek2-guide-F | CACCGACATCGTTCGTTACTATGAT |
| Nek2-guide-R | aaacATCATAGTAACGAACGATGTC |
| Rootletin-guide-F | CACCGAATGGCGAGCTCATCGCGCT |
| Rootletin-guide-R | aaacAGCGCGATGAGCTCGCCATTC |
| Cnap1-guide-F | CACCGCGGCTGCAGAAGCTCACTG |
| Cnap1-guide-R | aaacCAGTGAGCTTCTGCAGCCGC |

**Supplementary Table 3:** Oligo sequences for shRNA cloning into Tet-pLKO

| Name | Sequence (5' - 3') |
|---|---|
| shKIF2C-CDS-F | CCGGGCCCGAATGATTAAAGAATTTCTCGAGAAATTCTTTAATCATTCGGGCTTTTTG |
| shKIF2C-CDS-R | AATTCAAAAAGCCCGAATGATTAAAGAATTTCTCGAGAAATTCTTTAATCATTCGGG |

**Supplementary Table 5:** The list of siRNA products used in the study

| Gene | Brand | Product No |
|---|---|---|
| Nek2A | MERCK | EHU109951 |
| Gas2L1 | MERCK | EHU078871 |
| TRF1 | MERCK | EHU114821 |
| NuMA | MERCK | EHU059141 |
| KIFC1 | MERCK | EHU148011 |
| KIF2C | MERCK | EHU046211 |

**Supplementary Table 6:** RT-qPCR primers used in this study

| Name | Sequence (5' - 3') |
|---|---|
| NEK2-F | TTG GAG CAG AAA GAA CAG GAG C |
| NEK2-R | TCC CCA CTG AAA TGA ACT TTC TTC |
| ACTB-F | TCA CCA TGG ATG ATG ATA TCG C |
| ACTB-R | ATA GGA ATC CTT CTG ACC CAT GC |
| GAPDH-F | CTG ACT TCA ACA GCG ACA CC |
| GAPDH-R | GTT GTC ATA CCA GGA AAT GAG C |
| PLK4-F | GGC CAA GGA CCT TAT TCA CCA |
| PLK4-R | TGT GGC ATG CCC ACT ATC AA |
| TERF1-F | CAG CGC AGA GGC TAT TAT TCA TGG |
| TERF1-R | AGG GCT GAT TCC AAG GGT GT |
| GAS2L1-F | GAC ACG CTG GAG CAT TAC CT |
| GAS2L1-R | TGG AGA AAA GGT GCA GAC CC |
| HSET-F | TTG GTA CTG CTC AGG CCA AC |
| HSET-R | GAT AGC CCT GGG ACA TGG TG |
| NuMa-F | CAG GTG GAA ACT AAT TCT AAG CCA G |
| NuMa-R | GTC ACT CCA ATG CGC CTC CT |
| KIF2C-F | CGC GTT TCT CTT CCT TGC TG |
| KIF2C-R | CCT TTG TGG CAC CTC CTT CT |
