## supplemental figures for "Less is More: Nek2A Unclusters Extra Centrosomes and Induces Cell Death in Cancer Cells via KIF2C Interaction"

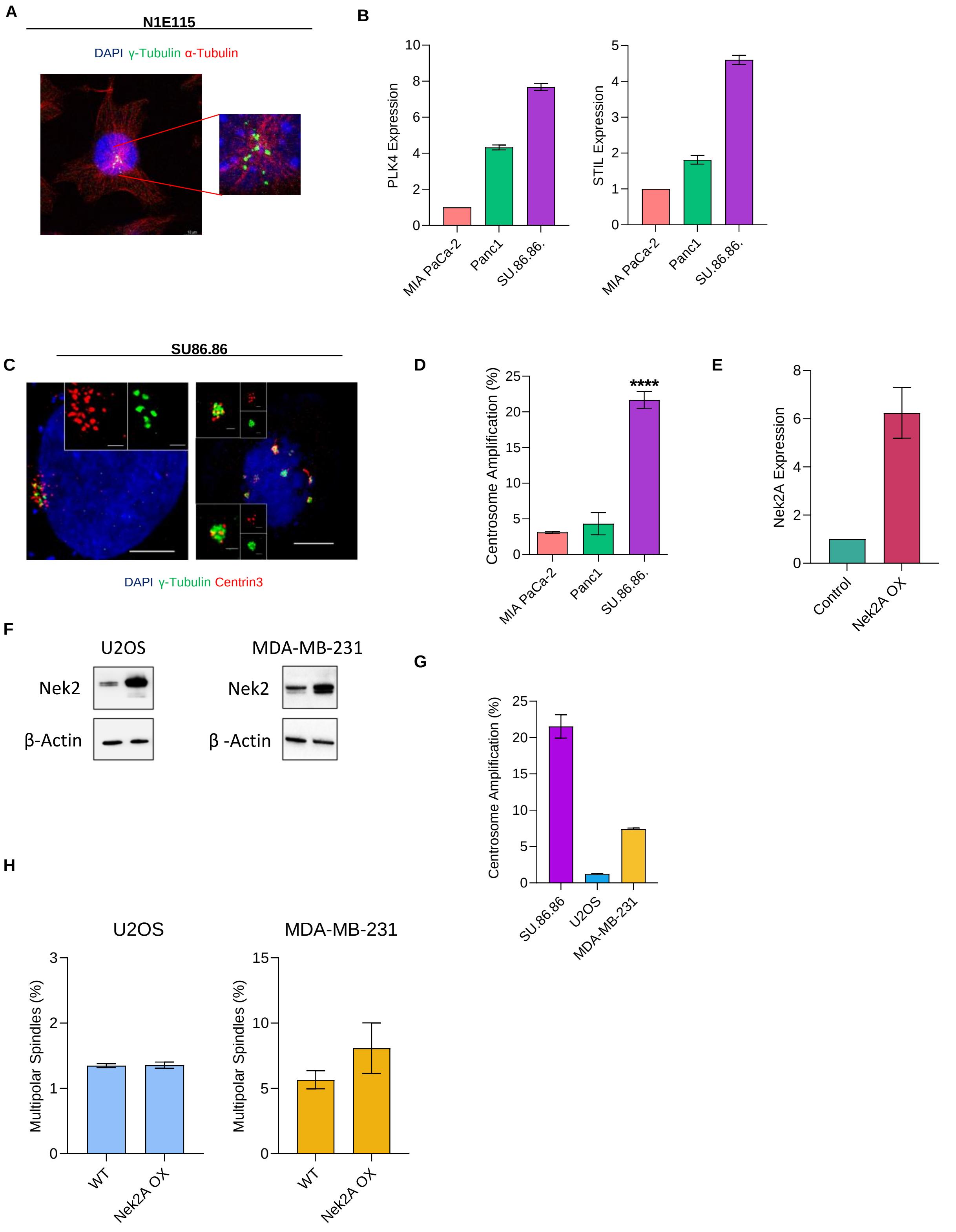


**Supplementary Figure 1:** (**A**) Confocal microscopy image showing the presence of supernumerary centrosomes in N1E-115 cell line. Centrosomes were stained via anti γ-tubulin antibody and Alexa Fluor 488. (**B**) RT-qPCR analysis to verify mRNA expression levels of PLK4 and STIL in PDAC cell lines. Bars show the expressions relative to MIA PaCa-2 (normalized to 1) (**C**) Confocal microscopy image showing the presence of supernumerary centrosomes (γ-tubulin) and centrioles (Centrin3) in SU86.86 cell line. (**D**) Quantification of centrosome amplification levels (>2 centrosomes per cell) in PDAC cell lines. (**E**) RT-qPCR analysis to verify Nek2 overexpression. (**F**) Western Blot verification of Nek2 overexpression in U2OS and MDA-MB-231 cell lines. (**G**) Quantification and comparison of CA in selected cell lines. (**H**) Scoring percentage multipolar metaphases observed in U2OS and MDA-MB-231 cells. Experiments were performed as two biological repeats. Error bars show standard deviations. Statistical significance was shown as * : $p<0.05$, **: $p<0.01$, ***: $p<0.001$, **** : $p<0.0001$. OX: overexpression

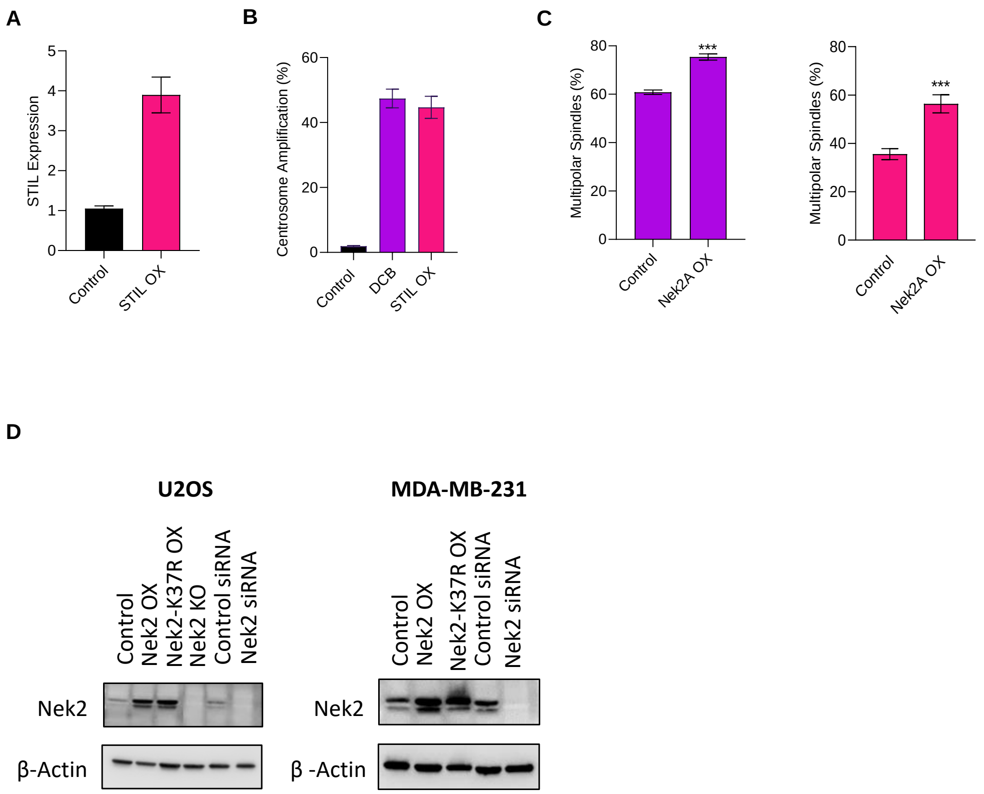


**Supplementary Figure 2:** (**A**) RT-qPCR analysis to verify STIL overexpression in U2OS cells. (**B**) Quantification of centrosome amplification levels (>2 centrosomes per cell) in DCB treated and STIL overexpressing U2OS cells. (**C**) Metaphase scoring results show that Nek2A OX induces significant increase in MPS formation in DCB (left) and STILL (right) CA models in U2OS cells. (**D**) Western Blot showing the protein levels of Nek2 in different experimental groups of U2OS and MDA-MB-231 cells. Experiments were performed as two biological repeats. Error bars show standard deviations. Statistical significance was shown as * : p<0.05, **: p<0.01, ***: p<0.001, **** : p<0.0001. OX: overexpression, Nek2-K37R: kinase-dead mutant, KO: knock-out

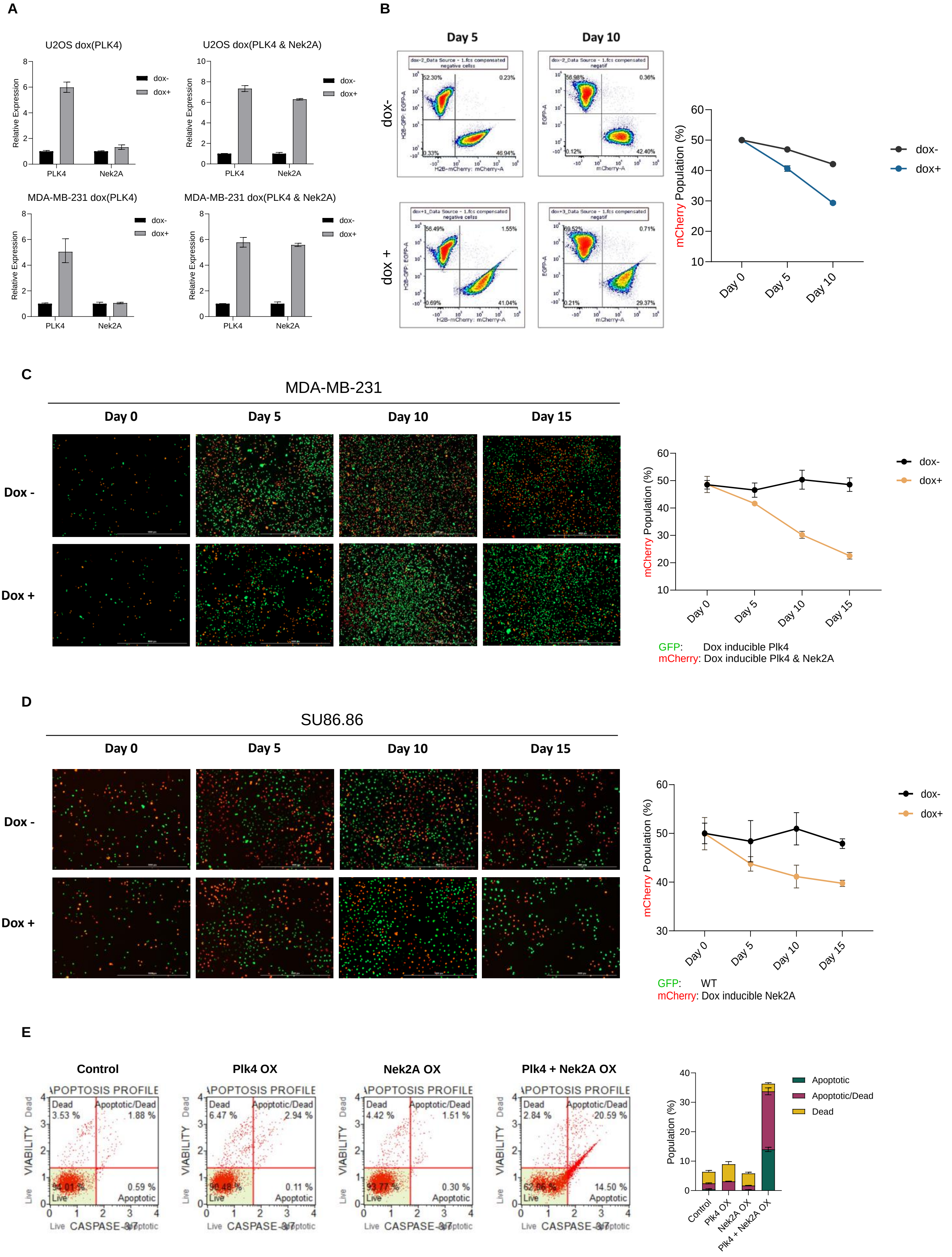


**Supplementary Figure 3:** (**A**) RT-qPCR analysis to verify dox-inducible PLK4 and Nek2 overexpression in U2OS and MDA-MB-231 cells. (**B**) Flow cytometry-based quantification of mCherry-tagged cell populations at day 0, 5 and 10 in U2OS cells. (**C**) Dual-colour competition assay and image-based quantification of mCherry-tagged Nek2 overexpressor MDA-MB-231 cells. Centrosome amplification was induced by PLK4 overexpression. (**D**) Dual-colour competition assay and image-based quantification of mCherry-tagged Nek2A overexpressor SU86.86 cells. (**E**) Caspase 3/7 assay confirming the apoptosis induced by multipolar metaphases in U2OS cells. Experiments were performed as two biological repeats. Error bars show standard deviations. Statistical significance was shown as * : p<0.05, **: p<0.01, ***: p<0.001, **** : p<0.0001. OX: overexperssion

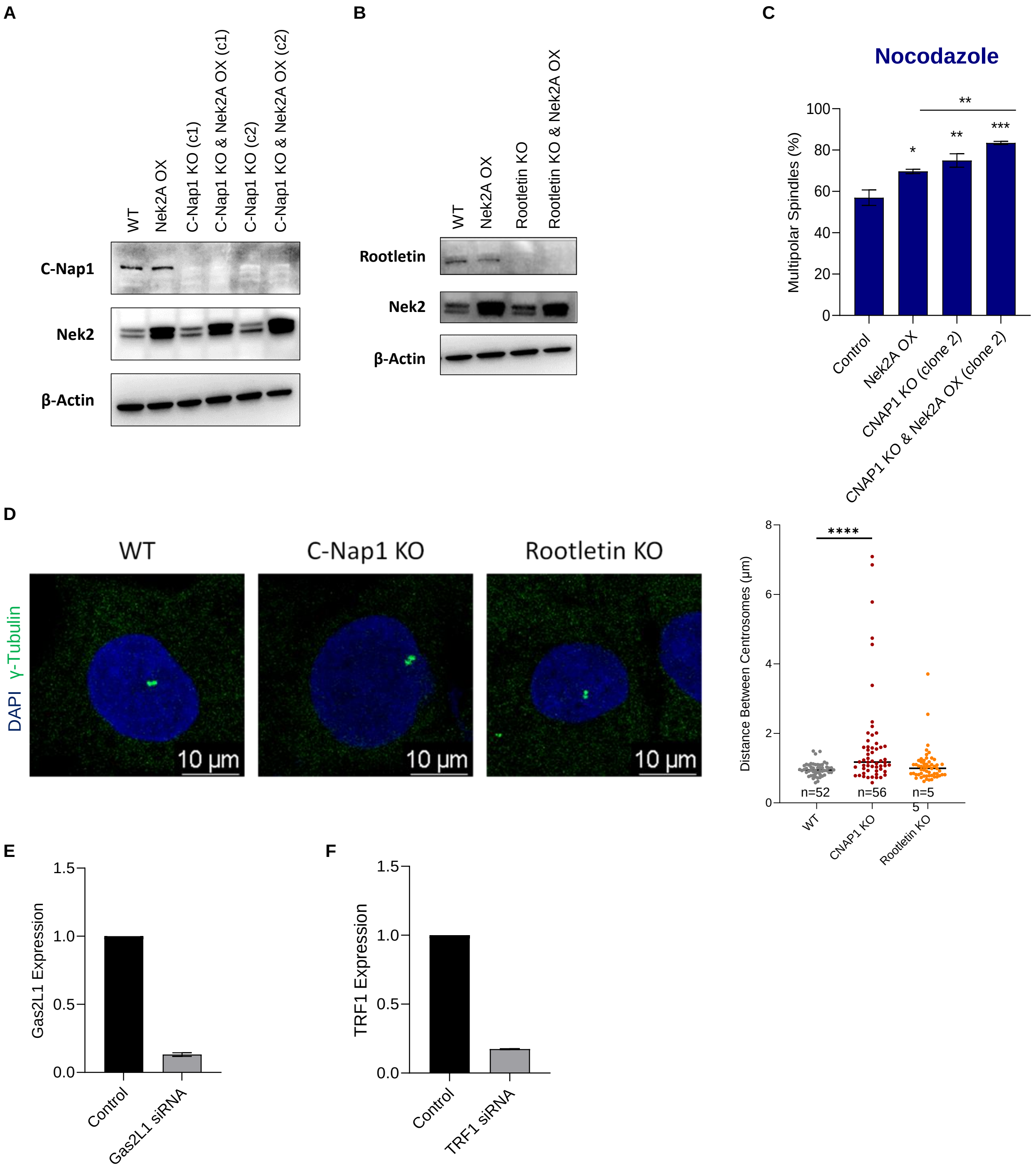


**Supplementary Figure 4:** (**A-B**) Western Blot verifications of CRISPR/Cas9 mediated KO of CNAP1 and Rootletin, along with exogenous Nek2 overexpression (**C**) Metaphase scoring of independent CNAP KO monoclone. (**D**) Immunofluorescence staining of centrosomes (γ-tubulin) to measure the distance between the centres of γ-tubulin foci by confocal microscopy. Graph indicates the median centrosome distances. **(E-F)** RT-qPCR analysis verifies siRNA mediated suppression of Gas2L1 and TRF1. Experiments were performed as two biological repeats. Error bars show standard deviations. Statistical significance was shown as * : p<0.05, **: p<0.01, ***: p<0.001, **** : p<0.0001. OX: overexpression, KO: knock-out

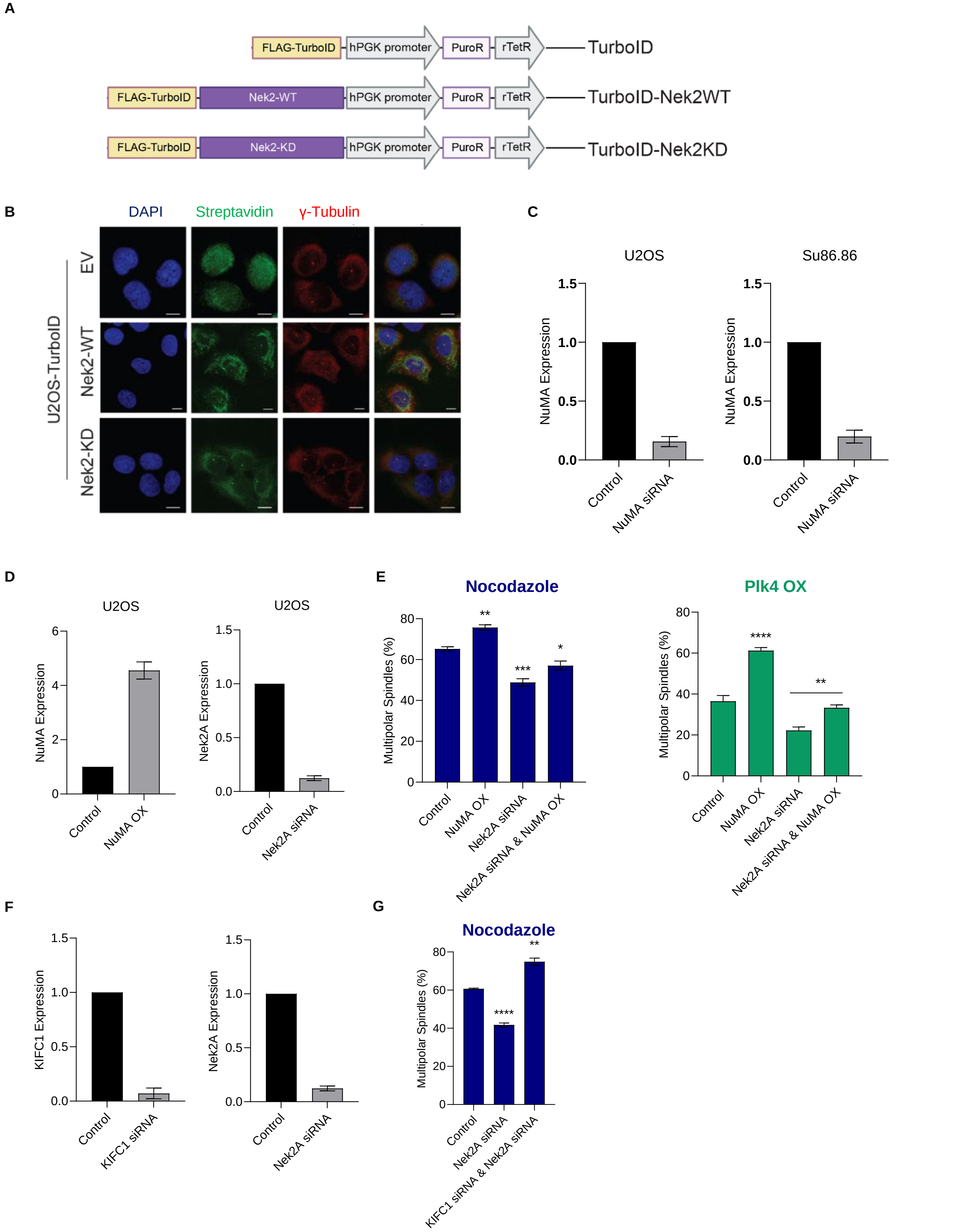


**Supplementary Figure 5:** (**A**) Turbo-ID proximity labelling system to identify interaction partners of Nek2A in U2OS cells. (**B**) Cellular localization of TurboID-Nek2A-WT and KD, verified by IF staining. (**C**) RT-qPCR analysis to verify siRNA-mediated knockdown of NuMA in U2OS and Su86.86 cells. (**D**) RT-qPCR analysis results to verify ectopic NuMA overexpression and siRNA-mediated Nek2A knockdown in U2OS. (**E**) Additional metaphase scoring experiments to assess the relationship between NuMA and Nek2A on centrosome clustering events in U2OS. (**F**) RT-qPCR analysis results to verify siRNA-mediated knockdowns of KIFC1 and Nek2A in U2OS cells. (**G**) Additional metaphase scoring experiment to examine the independence of KIFC1 on regulating centrosome clustering. Experiments were performed as two biological repeats. Error bars show standard deviations. Statistical significance was shown as * : p<0.05, **: p<0.01, ***: p<0.001, **** : p<0.0001. WT: wild type, KD: kinase-dead, OX: overexpression

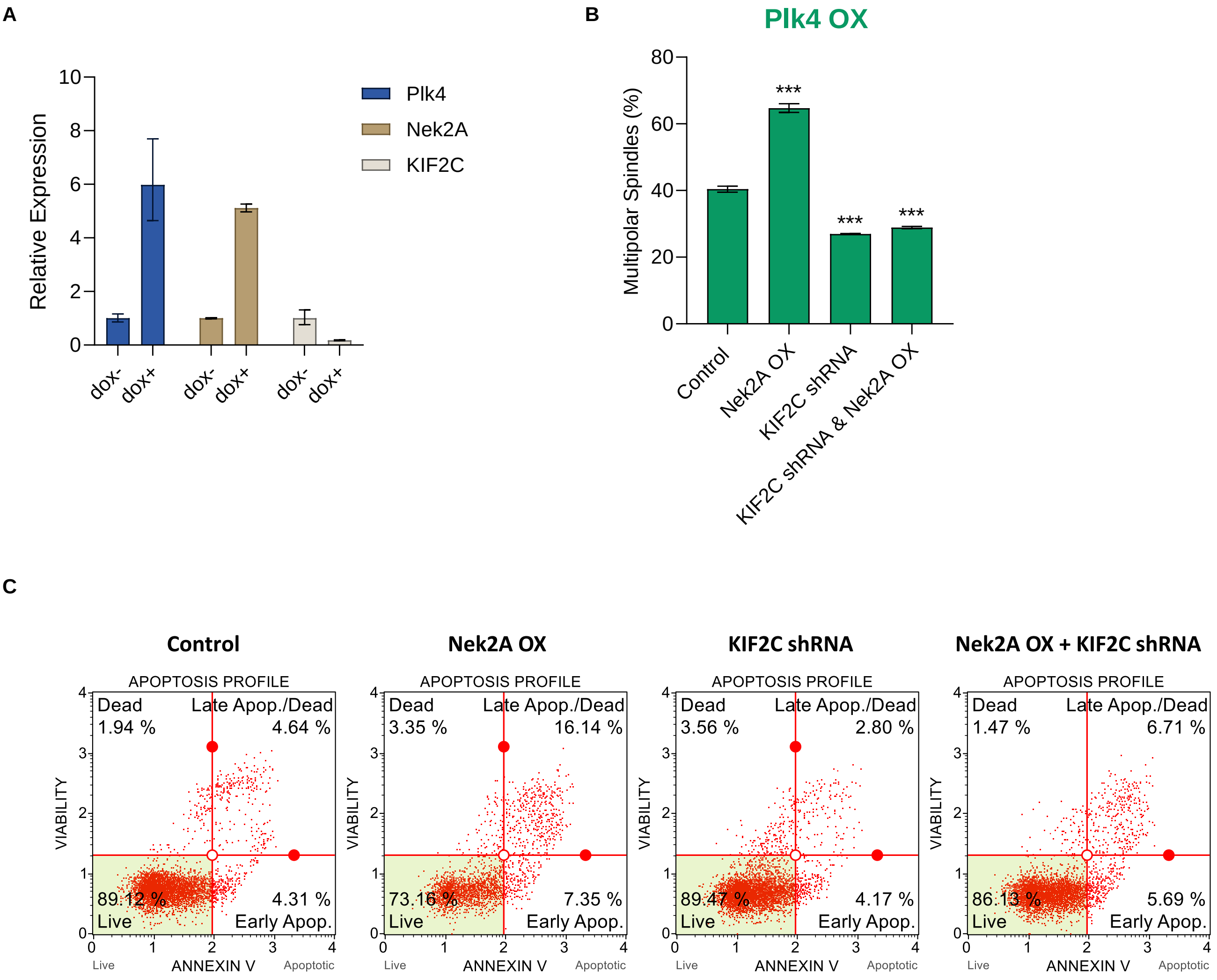


**Supplementary Figure 6:** (**A**) ) RT-qPCR analysis to verify dox-inducible overexpressions of Plk4 and Nek2A and dox-inducible shRNA of KIF2C. (**B**) Metaphase scoring data verifying the effect of KIF2C silencing on Nek2A's centrosomal un-clustering activity. (**C**) Representative plots of Annexin V apoptosis assay examining he importance of KIF2C for Nek2A to exert its activity. Experiments were performed as two biological repeats. Error bars show standard deviations. Statistical significance was shown as * : p<0.05, **: p<0.01, ***: p<0.001, **** : p<0.0001. OX: overexpression

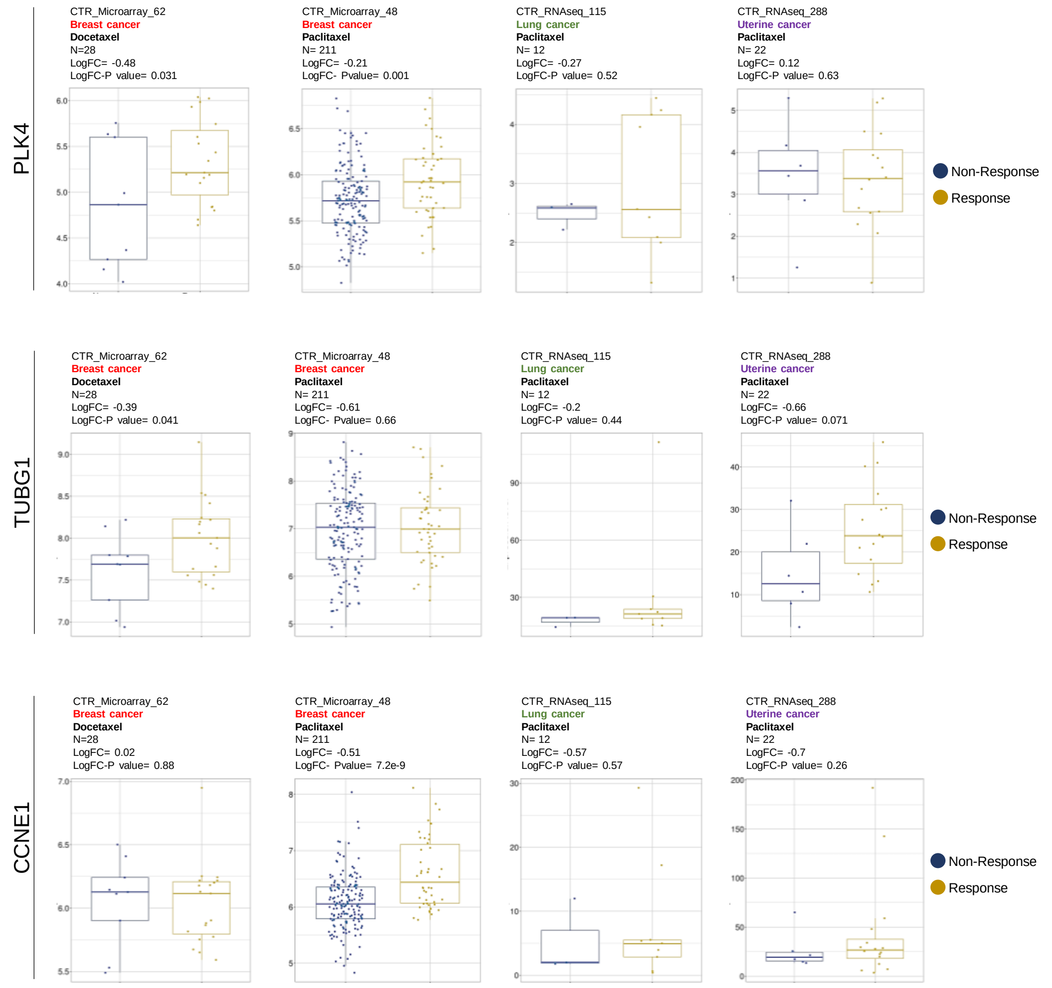


**Supplementary Figure 7:** Patient-derived clinical transcriptome data with taxane treatment response. In cohorts of cancer patients exhibiting positive response to treatment, relatively higher expression levels of centrosome amplification biomarkers (PLK4, TUBG1 and CCNE1) are observed compared to the non-responsive patients. Data was obtained from Cancer Treatment Response Gene Signature Data Base (CTR-DB).
